# Working memory adaptively recomputes plans amid distraction

**DOI:** 10.64898/2026.08.03.742569

**Authors:** Ziyao Zhang, Jarrod A. Lewis-Peacock

## Abstract

Navigating daily tasks requires working memory to retain information, formulate plans, and execute goal-directed actions. While frequent distractions may momentarily disrupt planned actions, individuals typically exhibit the cognitive resilience necessary to regroup and resume goal-directed behavior. How people adapt to these perturbations in dynamic environments remains unclear. To address this, we developed a working memory paradigm inspired by the arcade game *Snake*, in which participants navigated an agent to collect memorized targets while EEG and eye-tracking data were recorded. On each trial, participants encoded the location of one or two targets (apples), maintained them during a delay, and then guided a virtual agent (snake) to collect them. Critically, on 50% of trials, a secondary task introduced high-value novel targets that required immediate pursuit; these interruptions altered the agent’s spatial position, necessitating a re-evaluation of the original plan upon returning to the primary goal. Behaviorally, participants prioritized the target proximal to the agent and flexibly revised their collection order after distraction according to the agent’s updated position. Eye-movement analyses showed structured gaze sweeps during encoding that were consistent with efficient route planning. Representational similarity analyses of EEG data revealed that agent-centered object vectors were represented more strongly than absolute object locations during the delay. Following distraction, object vectors were recomputed relative to the agent’s new position, and representational strength was redistributed when the action plan was updated. Whereas neural representations were biased toward the distal, future-relevant target, gaze was preferentially directed toward the proximal, immediately relevant target, suggesting complementary roles for internal attention and external attention in multistep planning. During distraction, backward gaze sweeps revisited memorized target locations, indicating that replanning unfolded dynamically while the intervening task was still ongoing. These data demonstrate that prospective neural coding and attentional sampling coordinate to adapt behaviors to the shifting demands of goal pursuit in dynamic environments.

## Introduction

Adhering to the pursuit of goals in a busy, distracted world requires the ability to preserve intentions over time in order to guide behavior effectively. Working memory is a core cognitive function that supports the active maintenance and manipulation of task-relevant information for future use (Baddeley 2003; van Ede and Nobre 2023). In classic delay paradigms in the laboratory, a transient sensory stimulus provides information that must be internally maintained across a memory delay to guide a later response (Luck and Vogel 1997, 2013). Yet, for working memory to support flexible behavior outside of the laboratory, it must not only interact with other cognitive functions such as planning and decision-making, but also continuously interface with the external world, where new information must be integrated into ongoing actions. Despite their ecological importance, these functions have largely been studied in isolation, and it remains poorly understood how the brain coordinates working memory and planning in the service of goal-directed behavior.

Internal representations that are associated with working memory maintenance have been primarily studied using encoding and decoding models applied to neural data (Bettencourt and Xu 2016; Hallenbeck et al. 2021; Harrison and Tong 2009; Lewis-Peacock et al. 2012; Panichello and Buschman 2021; Rademaker, Chunharas, and Serences 2019; Serences et al. 2009; Sprague, Ester, and Serences 2016; Wolff et al. 2017). In humans, mounting evidence highlights the dominance of sensory codes in sustaining working memories (Hallenbeck et al. 2021; Harrison and Tong 2009; Lewis-Peacock et al. 2012; Serences et al. 2009). Decoding models can reliably predict the category of visual information from neural responses for both low-level features such as spatial locations, colors, and orientations, as well as higher-level image categories. This body of work aligns with the sensory recruitment hypothesis of working memory, which posits that sensory regions maintain sensory specific representations to support memory (D’Esposito 2007; D’Esposito and Postle 2015; Postle 2006, 2015). More recent studies, though, have revealed that working memory also involves widely distributed coding across regions such as the intraparietal sulcus (IPS) and frontal eye fields (FEF) (Christophel et al. 2017; Christophel, Hebart, and Haynes 2012; Curtis and Sprague 2021; Hallenbeck et al. 2021; Lorenc et al. 2018; Sprague et al. 2016). Yet, even these distributed representations seem to be primarily sensory in nature, with their distinctiveness arising largely from differences in the sensory features of the memoranda. This sensory-dominant view is partly driven by a potential confound in traditional working memory paradigms, where the sensory stimulus itself often serves as the primary task-relevant variable, directly supporting later recognition or reproduction.

An alternative account emphasizes the maintenance of prospective plans in addition to retrospective sensory information. Recordings of prefrontal delay activity have revealed correlations not only with sensory input, but also with diverse variables that are associated with action planning, expected stimuli, and abstract rules (Fascianelli et al. 2024; Rainer, Rao, and Miller 1999; Rigotti et al. 2013; Stokes et al. 2013; Wallis, Anderson, and Miller 2001).

Retrospective sensory input and prospective plans can be dissociated in task designs that decorrelate sensory, rule, and action information (Lewis-Peacock, Cohen, and Norman 2016; Lewis-Peacock and Postle 2008). For example, instead of requiring a pure sensory recognition or reproduction, participants may be instructed to respond with either the left or right hand depending on the sensory input (Boettcher et al. 2021; van Ede et al. 2019; Nasrawi, Boettcher, and van Ede 2023). In conditions where the final motor response can be preplanned (e.g., using the left hand), this motor plan can be reliably decoded from motor regions or motor-related beta oscillations (Boettcher et al., 2021; van Ede et al. 2019; Henderson, Rademaker, and Serences 2022; Nasrawi et al. 2023), while the strength of the sensory code decreased (Henderson et al. 2022). The emergence of motor codes when responses could be preplanned suggests that working memory is not merely the passive maintenance of sensory codes. Instead, working memory might be better characterized as the mapping from sensory information to future behavior through the interaction with planning.

However, working memory constantly faces the challenge of distractions that momentarily disrupt planned actions, especially in dynamic environments where disruptions are frequent and unpredictable (Bettencourt and Xu 2016; Clapp and Gazzaley 2012; Fukuda et al. 2022; Gazzaley and Rosen 2016; Hallenbeck et al. 2021; Lorenc, Mallett, and Lewis-Peacock 2021; Zhang and Lewis-Peacock 2023b, 2023a, 2025). When an unexpected change occurs in the environment, the agent must remap retrospective sensory information onto new action sequences to achieve the goal (Gresch et al. 2025). If working memory relies too rigidly on prospective plans while discarding retrospective sensory codes, such remapping might become difficult.

Conversely, retaining retrospective sensory information may provide a flexible basis for replanning by supporting new mappings between past sensory inputs and future responses, although these representations alone may be insufficient to specify the actions required. Thus, the coordination between retrospective and prospective memory codes may be critical for flexible goal-directed behavior, especially in dynamic environments where preplanned actions can be disrupted by changing environmental affordances.

To investigate how working memory supports resilient behaviors to changing demands imposed by the environment, we developed a dynamic working memory paradigm based on the classic arcade game “Snake” in which participants memorize the locations of target apples and then navigate a snake agent to collect them (Zhang and Lewis-Peacock 2026). In this task, participants consistently show a strong proximity bias, tending to collect apples from nearest to farthest, indicating that they configure efficient prospective action plans during encoding.

Critically, these initial plans were not static; an intervening task acted as a distraction requiring participants to ‘pin’ their primary goal, pursue a novel subgoal, and then dynamically recalculate their return to the original targets from the agent’s new position. This design allowed us to test how prospective plans would be adapted following distraction. Rather than rigidly adhering to initial plans, participants demonstrated adaptive flexibility. They dynamically updated their priorities of target collections based on their locations relative to the agent’s post-distraction coordinates. In the current study, we sought to clarify the mechanisms that support such flexible replanning in working memory. To do so, we collected EEG and eye-tracking data as participants completed the game.

Two central questions we aimed to address were what representations support the formation of action plans in working memory, and how do these representations enable flexible adaptation to changes in the environment? Our main hypothesis was that the coordination of retrospective sensory information and prospective plans support flexible replanning in response to disruptions. We operationalized retrospective sensory information and prospective plans in our task based on the relevant spatial navigation and cognitive map literatures (Behrens et al. 2018; Epstein et al. 2017; Høydal et al. 2019; Ormond and O’Keefe 2022; Tang et al. 2026; Yu et al. 2026). Accordingly, retrospective sensory information is operationalized as the *object information* available during encoding. In the context of our paradigm, this refers to the coordinates of the apple targets. By contrast, prospective plans are operationalized as *object vectors;* in this task, they represent the coordinates of each apple target relative to the snake agent. These vectors can capture potential actions more directly than absolute object coordinates, as they correspond to the sequence of movements required to collect the apples (Høydal et al. 2019; Ormond and O’Keefe 2022; Tang et al. 2026; Yu et al. 2026). In addition, we defined a *prospective object vector*, which captures the spatial relations between targets that can inform a future step in a multistep plan. Retrospective sensory information and prospective plans are often intertwined in classical spatial working memory paradigms, where the remembered location is both the information encoded into working memory and the critical component of the prospective plan guiding a saccade moving from central fixations toward that location. Our paradigm enables us to dissociate the two representations by introducing the dynamic agent state.

Specifically, retrospective sensory information and prospective plans can both be encoded in our task, when all targets and the agent location are simultaneously visible. However, they can be dissociable after the secondary distraction task. When the snake’s position is shifted, the object information (original absolute coordinates of the targets) remains unchanged, whereas the object vectors (relative coordinates of the targets to the agent’s new position) would need to be updated in order to accurately guide collection of the targets. Prospective plans are particularly informative for our task because they contain information about potential actions needed to reach target locations, as well as plans for multistep action sequences. However, relying on prospective plans alone would be insufficient when unexpected environmental interruptions occur and those plans become outdated. We therefore hypothesized that the coordination between retrospective sensory information and prospective plans guides flexible replanning in working memory.

## Methods

### Participants

25 participants were recruited through SONA at the University of Texas at Austin (Age: M = 19.12, SD = 1.20; 16 females, 9 males). Participants provided consent via REDCap (https://www.project-redcap.org/) and received course credit as compensation. The study was approved by the institutional review board. Eye-tracking data from one participant was not collected due to technical issues, leaving a final sample of 24 participants for eye-tracking analyses.

### Stimuli

The apple, bush, star, and snake images were downloaded from Flaticon, an online platform that provides freely accessible visual icons (https://www.flaticon.com). Sound stimuli, including the trial start sound, background music, reward sound, distraction music, and penalty sound, were obtained from Pixabay, an online platform that provides free sounds (https://pixabay.com/). All study materials are available on OSF. The game, inspired by the classic *‘Snake’* arcade game, was developed in Python using Pygame (https://www.pygame.org), and the code for generating the game display is shared.

### Experimental procedure

Each trial began with a 2s encoding phase, during which 16 green cartoon bushes (2.2° × 2.2°) were displayed across four quadrants of the screen (four per quadrant), with a minimum spacing of 2.2° between bushes. Either 1 or 2 red cartoon apples (2.2°) appeared on randomly selected bushes. A blue cartoon snake (head: 2.2°; body: 2.2° × 4.4°, RGB: 75, 179, 253) was presented at the center of the screen, initially facing right in the first trial of each game session. Participants were instructed to memorize the apple locations and later control the snake to collect all presented apples after they disappeared (**Fig 1a & Video 1**).

**Fig. 1.**
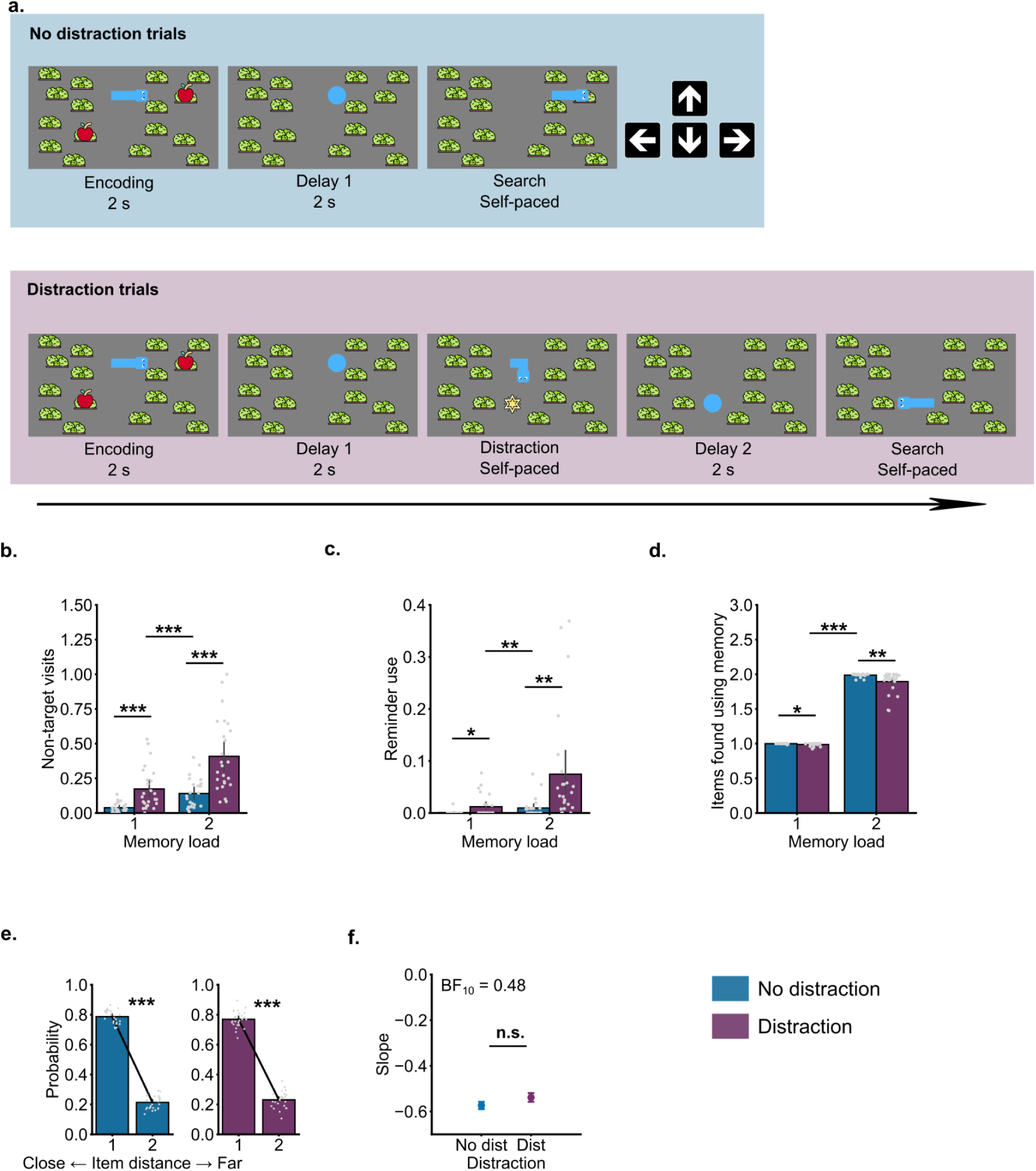
Task illustration and behavioral proximity bias. a). Participants encoded the locations of one or two target apples and then collected them after a delay, during which they might be required to complete an additional distracting task of collecting the star item. b-d). Participants’ behavioral errors were measured by visits to non-target locations and reminder use, as well as by the estimated number of items collected using memory. e). Probability of an item being collected as the first target as a function of its relative distance to the agent’s position. f). Slopes of linear models fitted to the probability of an item being collected first as a function of distance. n.s. indicates p > .05; * indicates p < .05; ** indicates p < .01; *** indicates p < .001. Error bars represent 95% confidence intervals. [snakeegg_examples.mp4](https://drive.google.com/file/d/1xLtw1p3IDYvKkC2bFukmONLqC4i04yLI/view?usp=sharing) **Video. 1. Task illustration.**

At the end of the encoding phase, both the apples and the snake disappeared, while the bushes remained visible and stayed in the same spatial locations. A blue circle (RGB: 75, 179, 253) appeared at the snake’s head position for 2s, serving as a fixation point. Participants were instructed to fixate on the blue circle, minimize blinks, and avoid eye movements during this delay phase. The circle (initially 2.2° in diameter) gradually shrank as time went by and disappeared at the end of the delay phase. In no-distraction trials (50%), the snake then reappeared and began moving at a constant speed of 6°/s. Participants controlled its movement using the direction buttons (up, down, left, and right) on a gamepad controller.

An apple was collected when the snake reached the corresponding bush. Successful collection triggered visual and auditory feedback: the apple reappeared at its original location for 0.5s, accompanied by a sound cue and a green “+1” next to the snake, indicating a point gain. If the snake visited a non-target bush, an apple core image appeared at the location, an error sound was played, and the snake was frozen for 1s as a penalty. A red “-1” was displayed next to the snake to indicate a point deduction.

In distraction trials (50%), after the delay phase, a star image appeared at a randomly selected location on screen, at least 2.2° away from any bush. The distance between the star and apple targets were also randomized. Participants were instructed to collect the star before retrieving the apples. The star was worth 3 points, incentivizing participants to prioritize it during the distraction phase. Importantly, the star disappeared if an apple was collected first, ensuring that participants had to complete the distraction task before returning to the main task of finding apples. For our analyses, we focused on distraction trials where participants collected the star before collecting any apples. Once the star was collected and the distraction phase ended, a second delay phase occurred, during which a gradually shrinking blue circle reappeared at the snake’s head position for 2s. Participants were again instructed to fixate on the blue circle, minimize blinks, and avoid eye movements. The snake reappeared following the second delay, and participants proceeded to collect the apples.

If participants were uncertain about the apple locations, they could activate a reminder function by pressing the ‘x’ key on the gamepad controller. This briefly revealed all uncollected apples for 2s, allowing for re-encoding. However, using the reminder incurred a 1-point penalty and introduced an additional 2s freeze penalty, during which the snake agent was replaced by the blue fixation circle that participants had to fixate before resuming movement.

A trial ended once all the apples were collected. The bushes were then removed, and the snake automatically moved along a predefined square path, creating a brief dancing animation. The next trial began with a new arrangement of bushes and apples, while the snake’s position carried over from the previous trial. This design prevented participants from learning stable spatial configurations of potential target locations and instead required them to rely on working memory to complete the task. An experimental run ended if the snake collided with the screen boundary, automatically triggering the start of a new run. The experiment ended when participants completed five sessions of gameplay, with each session lasting approximately 20 minutes. On average, participants completed 335 trials (SD = 36.3 trials).

### Eye-tracking acquisition

Eye movements were recorded using GazePointer GP3, with a sampling rate of 60 Hz. Participants were seated approximately 100 cm from the monitor (1920 × 1080 px) in a head-stabilized chin rest to minimize motion artifacts. Before each session, a 9-point calibration procedure was performed to ensure spatial accuracy (mean error < 0.5° visual angle). During calibration, participants sequentially fixated on nine predefined locations arranged in a 3 × 3 grid: the center of the screen, the four corners, and the midpoint of each screen edge. Raw gaze coordinates were saved and used for analyses.

### EEG acquisition

The EEG signal was acquired from a 64-channel Biosemi ActiveTwo system laid out according to the extended international 10–20 system. Data was recorded at 1,024 Hz. Additional electrodes were placed on the mastoids, below the right eye, to the left of the left eye, and to the right of the right eye.

### EEG preprocessing

The data was re-referenced to the average of mastoids, down-sampled to 500 Hz, and bandpass filtered (0.1 Hz high-pass and 40 Hz low-pass) using functions from an open source Python module, MNE (Gramfort et al. 2013). Two sets of epochs were created, relative to the onset of the first delay window following the encoding phase, and relative to the onset of the second delay window following the distraction phase in distraction trials. Artifact detection and trial rejection were performed for both sets of epochs during the delay windows using an automated artifact rejection algorithm, Autoreject, implemented in Python (Jas et al. 2017). Epochs were visually inspected following automated rejections to exclude additional trials that contained artifacts associated with excessive eye movement, blink, and motion activities. For the participants used in analyses, we rejected on average 3.7% of trials (SD = 8.4% of trials).

### Decision biases

To assess proximity biases in participants’ actions, we ranked target locations relative to the agent’s (snake) position in each trial and calculated the probability of collecting the first item based on its distance. If participants prioritized proximal targets, the likelihood of collecting an item first in a trial should decrease as its distance from the agent increases.

We used mixed linear models for statistical analyses, implemented with the brms package in R (Bürkner 2021). The ranked item distance and distraction were predictors, with the probability of an item being collected first as the outcome variable. For linear model results, we report β, 95% confidence intervals and Bayes factors.

### Gaze trajectory analysis

Regions of interest (ROIs) were defined as circles (d = 3°) centered on the agent location (snake head) and the target locations (apples). At each frame, gaze points were classified into four categories: the agent location (1), the proximal target (2), the distal target (3), or other locations (0). Gaze patterns were identified based on these classifications. An optimal forward sweep was defined as gaze transitions from the agent location to the proximal target and then to the distal target (e.g. 1-1-1-2-2-3-3-3). An optimal backward sweep was defined as transitions from the distal target to the proximal target and then back to the agent location (e.g. 3-3-2-2-2-2-1-1). A suboptimal forward sweep was defined as transitions from the agent location to the distal target and then to the proximal target (e.g. 1-1-1-3-3-2-2-2), while a suboptimal backward sweep was defined as transitions from the proximal target to the distal target and then back to the agent location (e.g. 2-2-3-3-3-3-1-1). In addition, independent encoding patterns were defined as gaze transitions from the agent location to either the proximal or the distal target and then back to the agent location (e.g. 1-1-2-2-1-1 or 1-1-3-3-1-1). We excluded gazes that landed on other locations to ensure robustness to blinks, to systematic eye-tracking errors and biases, and to drifts. For each pattern, the rate of occurrence was calculated as the number of occurrences relative to all identified gaze transitions.

This analysis was conducted during the encoding phase, when the target apples were visible on the screen, and during the distraction phase, when an additional star appeared to divert participants from collecting the apples. This allowed us to examine eye movement trajectories that are relevant for planning and replanning. The key difference between these phases was that, during the distraction window, the snake moved toward the star location. If replanning occurred, the updated action sequence should be anchored to the star location. We therefore conducted the gaze trajectory analysis relative to the star position during the distraction phase. By contrast, during encoding, the analysis was performed relative to the snake position, which remained fixed throughout the encoding window.

### Trial-level GLMM linking gaze patterns to proximity bias

We fitted generalized linear mixed-effects models (GLMMs) to test whether the frequency of each gaze pattern predicted whether participants ultimately selected the optimal target order on a given trial. Predictors were z-scored and then included in the model.

### Proportions of fixations

We used the same ROIs as in the gaze trajectory analysis. To estimate fixation patterns, we computed the proportion of fixations landing on the target locations relative to all valid fixations, using a moving window approach to ensure sufficient samples per window. Each window was 300 ms in length and moved in 50 ms steps.

Incidental eye movements, such as drifts or systematic biases, tended to land more often on the proximal target than the distal one when participants fixated on the agent location. To control for this distance effect, we randomly sampled locations that were the same distance as the two target locations relative to the agent locations. The logic is that if fixation proportions are determined primarily by physical distance, then those randomly sampled locations should have the same proportion of fixations as the actual targets. Any additional fixations on the true target locations, relative to the distance-matched controls, would therefore reflect voluntary eye movements guided by memory. For each trial, we randomly sampled control locations 100 times based on distance of the two target locations to the agent location, to establish an empirical null distribution. The final proportions of fixations on the targets were then baseline-corrected relative to these distance-controlled locations.

### Eye-tracking control analyses for sampling-rate variability

During the analysis of the eye-tracking data, we found that the sampling rate progressively decreased over the course of each recording session (M = 34.66 hz, SD = 21.46). This issue likely arose because eye movements were recorded on the same computer that displayed the experimental program. This potentially overloaded the system’s computational resources. Given the substantial variability in sampling rate, we conducted additional control analyses to determine whether our main eye-tracking findings were sensitive to these fluctuations. Specifically, for each participant, we resampled the eye-tracking data to the lowest trial-level empirical sampling rate and repeated the analyses using the resampled data.

### Trial-level representational similarity analysis (tRSA)

Our key analyses of EEG data were based on a trial-level variation of the representational similarity analysis (Huang et al. 2026; Kriegeskorte, Mur, and Bandettini 2008), which involved computing representational dissimilarity matrices (RDMs) based on hypothetical representational spaces and actual neural signals, and quantifying representation of each feature using the correlation approach. All analyses were implemented using customized functions in Python.

Model RDMs were computed for object locations and object vectors based on the Cartesian coordinates of the agent and the two targets in memory load 2 trials. Two object locations were extracted: the proximal target location (*x*_1_, *y*_1_), and the distal target location (*x*_2_, *y*_2_). Object vectors were defined as the spatial relations between target positions and the agent position (*x*_0_, *y*_0_). Specifically, the proximal object vector was (*x*_1_− *x*_0_, *y*_1_− *y*_0_), and the distal object vector was (*x*_2_− *x*_0_, *y*_2_− *y*_0_). In addition, we computed the prospective object vector, defined as the vector from the proximal to the distal target location (*x*_2_− *x*_1_, *y*_2_− *y*_1_). Each model RDM was constructed as a square matrix with dimensions equal to the number of trials of interest, where each cell reflected the Euclidean distance in the given spatial dimension between two trials. For example, to compute the dissimilarity between one trial where the proximal target was positioned at (x=200 pixels, y=100 pixels) and another trial at (x=500 pixels, y=500 pixels), we calculated the Euclidean distance: (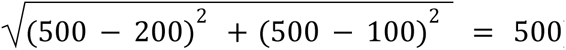). This value represented the dissimilarity between the two trials in the proximal object model RDM. In total, five model RDMs were generated (see **SFig 3**), each capturing the representational similarity structure expected if the brain encoded one of these spatial features or relations.

To construct the neural RDMs, we first z-scored the EEG voltage amplitudes at each channel across all conditions. At each time point, we extracted the 64-channel activation vector. We then computed one minus the Pearson correlation coefficient (1 - r) between pairs of trial-specific vectors, yielding an EEG pattern-based RDM with dimensions corresponding to the number of trials of interest.

To assess whether participants relied on object locations and/or object vectors to complete the task, we computed the representational similarity between the EEG RDM and the five model RDMs described above. We applied rank-based Spearman correlation analysis. The rank-based Spearman correlation was chosen because it is robust to non-linear relationships and scale differences across RDMs. Rather than correlating full neural and model RDMs at the participant level, we implemented a trial-wise approach. Specifically, for each trial, we extracted the vector of neural dissimilarities between that trial and all remaining trials. This vector constituted the neural representational profile for that trial at that time point. The corresponding trial-specific vector was then extracted from each model RDM. Trial-level RSA strength was defined as the correlation between the neural dissimilarity profile and the model dissimilarity profile. This procedure yielded, for each participant, a time-resolved RSA estimate for every trial and every model.

We used permutation-based tests to assess the statistical significance of group-level representational similarity for each spatial feature. For each of the five model RDMs, we extracted the unique dissimilarity values from the upper triangle and permuted the corresponding trial indices 1000 times to generate empirical null RDMs. This shuffling approach randomized trial structures while preserving trial-wise correspondence across RDMs. For each permutation, we computed the correlation between the EEG RDM and the model RDMs. The resulting correlations were averaged across participants to construct a group-level null distribution for each spatial feature.

We then compared the true group-level correlation derived from the true RDMs with its corresponding null distribution. A feature was considered significantly represented in EEG activation patterns if the observed similarity exceeded the 99th percentile of the null distribution (one-sided test, *p* < 0.01).

The same approach was used in memory load 1 trials, with the only difference that there was one target location in those trials instead of two target locations.

### Trial-level GLMM linking representational strength to memory error and proximity bias

We fitted generalized linear mixed-effects models (GLMMs) to test whether the trial-level representational strength of object locations and object vectors predicted whether participants ultimately selected the optimal target order on a given trial and whether participants made memory errors by visiting non-target locations or used reminders. Predictors were z-scored and then included in the model.

### Statistics

We conducted repeated-measures ANOVA to test main effects of independent variables and the potential interactions between them. Paired-samples t-tests were used to follow up on pair condition differences. Sidak correction was applied for multiple comparison corrections. All analyses were performed in Python using its associated statistical packages. Bayes factors for the alternative hypothesis (*BF10*) were reported, with values between 1 and 3 indicating anecdotal evidence, values between 3 and 10 indicating moderate evidence and values larger than 10 indicating strong evidence (Jeffreys 1961). Linear models were performed in R using the brms package (Bürkner 2021), and we report regression coefficients (β), 95% credible intervals, and Bayes factors.

## Results

As in our prior study (Zhang and Lewis-Peacock 2026), we analyzed participants’ behavioral errors in the distraction and no-distraction conditions. Non-target visits were measured as the rate of visiting non-target locations within a trial. Reminder use was defined as the rate of using reminders within a trial. Items found using memory were defined as the number of items found before the first reminder was used in a trial. Repeated-measures ANOVAs were conducted with memory load (1 vs 2) and distraction (no-distraction vs distraction) as within-subject factors. For non-target visits (**Fig 1b**), there was a significant interaction between memory load and distraction [*F*(1, 24) = 16.54, *p* < .001, 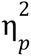 = .04, *BF10* = 6.03], along with main effects of memory load [*F*(1, 24) = 49.36, *p* < .001, 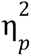= .22, *BF10* = 5. 6 × 10^6^] anddistraction [*F*(1, 24) = 50.42, *p* < .001, 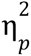= .29, *BF10* = 1. 4 × 10^9^]. For both load 1 and load 2 conditions, distracting events increased visits to non-target locations [load 1: *t*(24) = 5.29, *p* < .001, *d* = 1.24, *BF10* = 1146.12; load 2, *t*(24) = 6.94, *p* < .001, *d* = 1.33, *BF10* = 10^4^], with the detrimental effect of distraction being stronger under higher memory load. For reminder use (**Fig 1c**), there was a significant interaction between memory load and distraction [*F*(1, 24) = 10.48, *p* = .004, 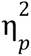 = .06, *BF10* = 5.51], as well as main effects of memory load [*F*(1, 24) = 10.69, *p* = .003, 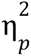= .09, *BF10* = 36.70] and distraction [*F*(1, 24) = 12.55, *p* = .002, 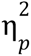 = .11, *BF10* = 75.53]. Distraction increased reminder use in both the load 1 condition [*t*(24) = 2.92, *p* = .015, *d* = 0.74, *BF10* = 6.06] and the load 2 condition [*t*(24) = 3.45, *p* = .004, *d* = 0.83, *BF10* = 18.38], with a stronger effect under load 2. For numbers of items found using memory (**Fig 1d**), there was a significant interaction [*F*(1, 24) = 12.05, *p* < .001, 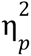 = .07, *BF10* = 9.26], along with main effects of memory load [*F*(1, 24) = 3762.63, *p* < .001, 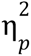= .97, *BF10* = 7. 1 × 10^77^] and distraction [*F*(1, 24) = 12.65, *p* < .001, 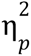 = .11, *BF10* = 54.05]. Distraction reduced the number of items found using memory in both the load 1 condition [*t*(24) = -2.92, *p* = .015, *d* = 0.74, *BF10* = 6.06] and the load 2 condition [*t*(24) = -3.54, *p* = .003, *d* = 0.85, *BF10* = 22.01], with a stronger effect under load 2. Replicating our prior findings, distraction impaired memory performance in the game, and this impairment was greater when memory load was higher.

### Updating of proximity biases following distractions

Our main behavioral analyses focused on participants’ planning behavior, which can be inferred from their collection order when multiple targets are present. We therefore restricted the following analyses to trials in the memory load 2 condition. Replicating our prior finding (Zhang and Lewis-Peacock 2026), participants flexibly updated their action plans (as reflected in collection order) to prioritize the newly proximal target after a distracting event occurred. In no-distraction trials, participants were significantly more likely to collect the proximal target before the distal one (**Fig 1c**. *β* = -0.57, *CI* = [-0.61, -0.54], *z* = -35.97, *p* < .001), reflecting a strong proximity bias consistent with efficient planning. In distraction trials, however, the pursuit of the additional distractor target forced the agent to move to a new location, likely rendering their initial plans suboptimal. We hypothesized that participants would flexibly update their actions and prioritize the newly proximal item. This was indeed the case: participants were again more likely to collect the updated proximal target before the distal one (**Fig 1e**. *β* = -0.54, *CI* = [-0.58, -0.50], *z* = -28.17, *p* < .001). Importantly, the magnitude of this proximity bias did not differ significantly between no-distraction and distraction trials (**Fig 1f**. *β* = -0.02, *CI* = [-0.18, 0.14], *z* = -0.21, *p* = .830, *BF10* = 0.48). We further examined distraction trials in which the optimal action plan should have been updated after the distracting event (68.3 % of distraction trials) versus trials in which the optimal action plan remained unchanged. In both trial types, we found comparable proximity bias (**SFig 1**. same optimal plan: *β* = -0.54, *CI* = [-0.60, -0.48], *z* = -17.35, *p* < .001; different optimal plan: *β* = -0.55, *CI* = [-0.59, -0.50], *z* = -22.74, *p* < .001;), with no significant difference between the two trial types (*β* = -0.003, *CI* = [-0.17, 0.16], *z* = -0.03, *p* = .974).

### Configuration of efficient plans during the encoding window

Research on goal-directed behaviors in naturalistic settings has shown that eye movements can reliably reflect internal planning (Ballard, Hayhoe, and Pelz 1995; Draschkow, Kallmayer, and Nobre 2021; Hayhoe and Ballard 2005; Keshava et al. 2024; Zhu et al. 2022). Two distinct gaze trajectory patterns have been associated with hierarchical planning: *forward sweeps*, where gaze shifts from the agent’s position toward intermediate transitions and final goals, and *backward sweeps*, where gaze moves from final goals back through transitions to the agent’s position (Lakshminarasimhan et al. 2020; Zhu et al. 2022).

Motivated by these findings, we examined whether participants exhibited similar gaze trajectories during the encoding window, which would indicate active configuration of efficient plans. The optimal action plan in our task was to move first toward the proximal target and then to the distal one, while the suboptimal plan would be to move first toward the distal target and then the proximal one. We therefore defined forward and backward sweeps for both optimal and suboptimal plans. For completeness, we also included gaze patterns associated with the independent encoding of the two targets (**Fig 2a&b**). A repeated-measures ANOVA revealed a main effect of gaze patterns, *F*(5, 115) = 13.58, *p* < .001, 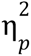 = .46, *BF10* = 4. 1 × 10^7^. Planned *t*-tests compared the occurrence rates of paired gaze patterns. Forward and backward sweeps aligned with optimal plans occurred more frequently than those aligned with suboptimal plans, although the difference was statistically significant only for forward sweeps. (**Fig 2c**, optimal forward vs. suboptimal forward: *t*(23) = 3.46, *p* = .002, *d* = 0.49, *BF10* = 18.30; optimal backward vs. suboptimal backward: *t*(23) = 1.93, *p* = .065, *d* = 0.32, *BF10* = 1.05). This pattern of eye gazes suggests that participants began configuring efficient action plans during the encoding window, which is consistent with the proximity bias we observed in their action sequences. Gaze patterns reflecting the independent encoding of individual items (e.g., agent → proximal target → agent) occurred more frequently for proximal than for distal targets, indicating that proximal targets were more likely to be encoded relative to the agent’s position, *t*(23) = 3.67, *p* = .001, *d* = 0.84, *BF10* = 28.25.

**Fig. 2.**
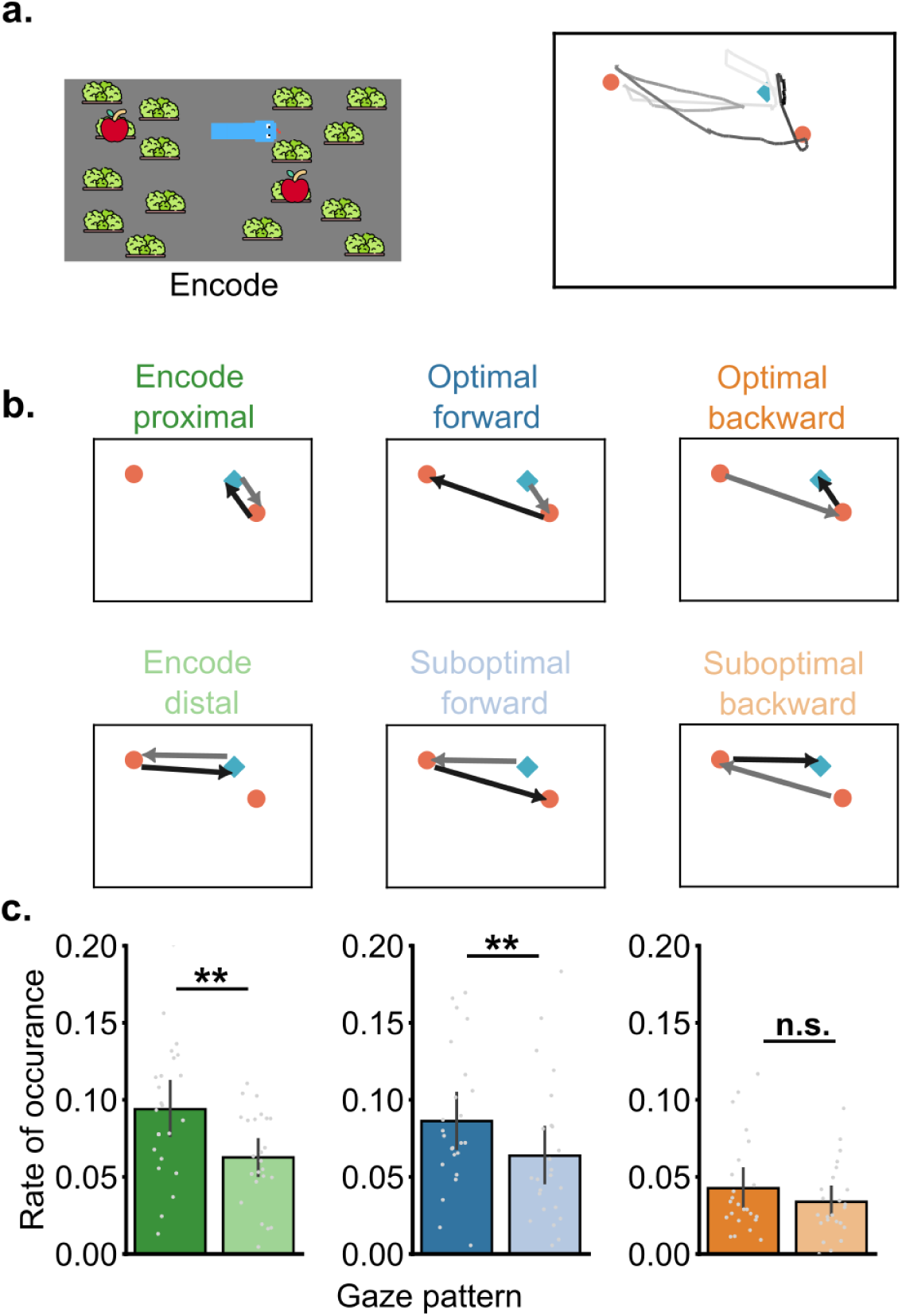
Gaze trajectories during the encoding window. a). Example gaze trajectories, with lighter colors indicating earlier time points and darker colors indicating later time points. b). Illustrations of gaze patterns in an example target display. c). Rate of occurrence for each gaze pattern. n.s indicates p > .05; ** indicates p < .01. Error bars represent 95% confidence intervals. [encode_combined.mp4](https://drive.google.com/file/d/1oYi5n5SeC7-SbLLph6op_8azVZSWfaGy/view?usp=sharing) **Video. 2.** Illustrations of gaze trajectories during the encoding and delay 1 periods from five example trials.

To examine the functional relevance of these gaze patterns, we used the rate of gaze patterns in a trial to predict whether that trial was solved using an optimal strategy (**SFig 2a**). Only gaze patterns associated with independent encoding of the proximal target positively predicted proximity bias (*β* = 0.12, *CI* = [0.01, 0.24], *BF10* = 1.59), whereas gaze patterns associated with independently encoding of the distal target negatively predicted proximity bias (*β* = -0.11, *CI* = [-0.20, -0.02], *BF10* = 1.83). We conducted control analyses and confirmed that these findings were robust to variability in the empirical sampling rate (**SFig 2b**).

### Maintenance of action plans through coordinated object location and object vector representations

Given the evidence for plan configuration during encoding, we next examined how action plans were maintained across the delay. We focused on two forms of spatial information: object locations and object vectors. Whereas object location representations preserve where the targets appeared, object vectors specify their spatial relations and can therefore provide more direct guidance for sequential action. However, because these vectors depend on the agent’s current position, they must be updated as the agent moves. To test how these spatial representations supported planning and replanning in working memory, we constructed hypothesized RDMs based on trial by trial differences in spatial information and tracked their neural representations throughout the delay periods (**SFig 3**). These RDMs capture the locations of the proximal and distal objects, the corresponding proximal and distal object vectors, and the prospective object vector representing the spatial relationship between the two targets. We hypothesized that participants would transform information held in working memory from retrospective object location representations to more action-relevant object vector representations. Critically, if participants maintained multistep plans, prospective object vectors capturing the spatial relationship between potential targets should also be represented during the delay.

We first compared the two candidate spatial representational frameworks in memory load 1 trials to test whether both object locations and object vectors were represented. During encoding and delay 1, object location and object vector representations coexisted with comparable strength. However, during the delay period, object vector representations were significantly stronger than object location representations, *t*(24) = 8.80, *p* < .001, *d* = 1.26, *BF10* = 2. 2 × 10 (**SFig 4a-b**). Following distraction, the updated object vector representation, defined as the vector connecting the updated agent location with the remembered target location, was stronger than both the object location representation and the initial object vector representation [updated object vector vs object location: *t*(24) = 8.98, *p* < .001, *d* = 1.36, *BF10* = 3. 2 × 10^6^; updated object vector vs initial object vector: *t*(24) = 9.85, *p* < .001, *d* = 2.57, *BF10* 1. 7 × 10^7^] (**SFig 4d-e**).

We next tested whether trial-level representational strength predicted memory errors across trials and participants while controlling for agent-target distance. In no-distraction trials, greater agent-target distance was associated with more visits to non-target locations (*β* = 0.30, *CI* = [0.00, 0.59], *BF10* = 2.52), whereas stronger object vector representations predicted fewer non-target visits (*β* = -0.81, *CI* = [-1.25, -0.29], *BF10* = 76.05, **SFig 4c**). A similar pattern emerged in distraction trials: greater agent-target distance was associated with more non-target visits (*β* = 0.33, *CI* = [0.14, 0.51], *BF10* = 104.63), whereas stronger updated object vector representations predicted fewer non-target visits (*β* = -0.36, *CI* = [-0.66, -0.07], *BF10* = 6.14, **SFig 4f**). Together, these findings indicate that stronger object vector coding was associated with fewer memory errors. Full results relating neural representational strength to memory errors in memory load 1 trials are reported in the Supplementary Materials.

We then tested whether object vector representations also dominated on memory load 2 trials and, critically, whether they were related to planned collection order. During encoding and delay 1, both object vector and object location representations were robust for the proximal and distal targets and persisted across both time windows (**Fig 3a&b**). Although all spatial representations were reliably represented (*p’s* <=.001), their mean correlation strengths differed significantly, *F*(4, 96) = 61.21, *p* < .001, 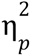 = .42, *BF10* = 2. 5 × 10 . Two main patterns emerged from the pairwise comparisons (**Fig 3c**). First, for both proximal and distal targets, object vector representations were stronger than object location representations, consistent with the results from memory load 1 trials [proximal object: *t*(24) = 7.43, *p* < .001, *d* = 1.86, *BF10* = 1. 1 × 10^6^; distal object: *t*(24) = 8.66, *p* < .001, *d* = 0.65, *BF10* = 1. 7 × 10^6^]. This pattern suggests a neural bias toward representing target information as spatial relations to the agent rather than retrospective object location coordinations. Second, the distal target was represented more strongly than the proximal target, regardless of representational format [object vector: *t*(24) = 7.75, *p* < .001, *d* = 1.32, *BF10* = 2. 6 × 10^5^; object location: *t*(24) = 7.81, *p* < .001, *d* = 2.03, *BF10* = 3. 0 × 10^5^]. This finding suggests a neural bias toward the distal, future target over the proximal, immediately relevant target. Finally, the prospective object vector was also reliably represented, *t*(24) = 8.08, *p* < .001, *d* = 1.62, *BF10* = 1. 3 × 10^8^. This vector was not directly relevant to the immediate action of moving from the agent’s current position to either target.

**Fig. 3.**
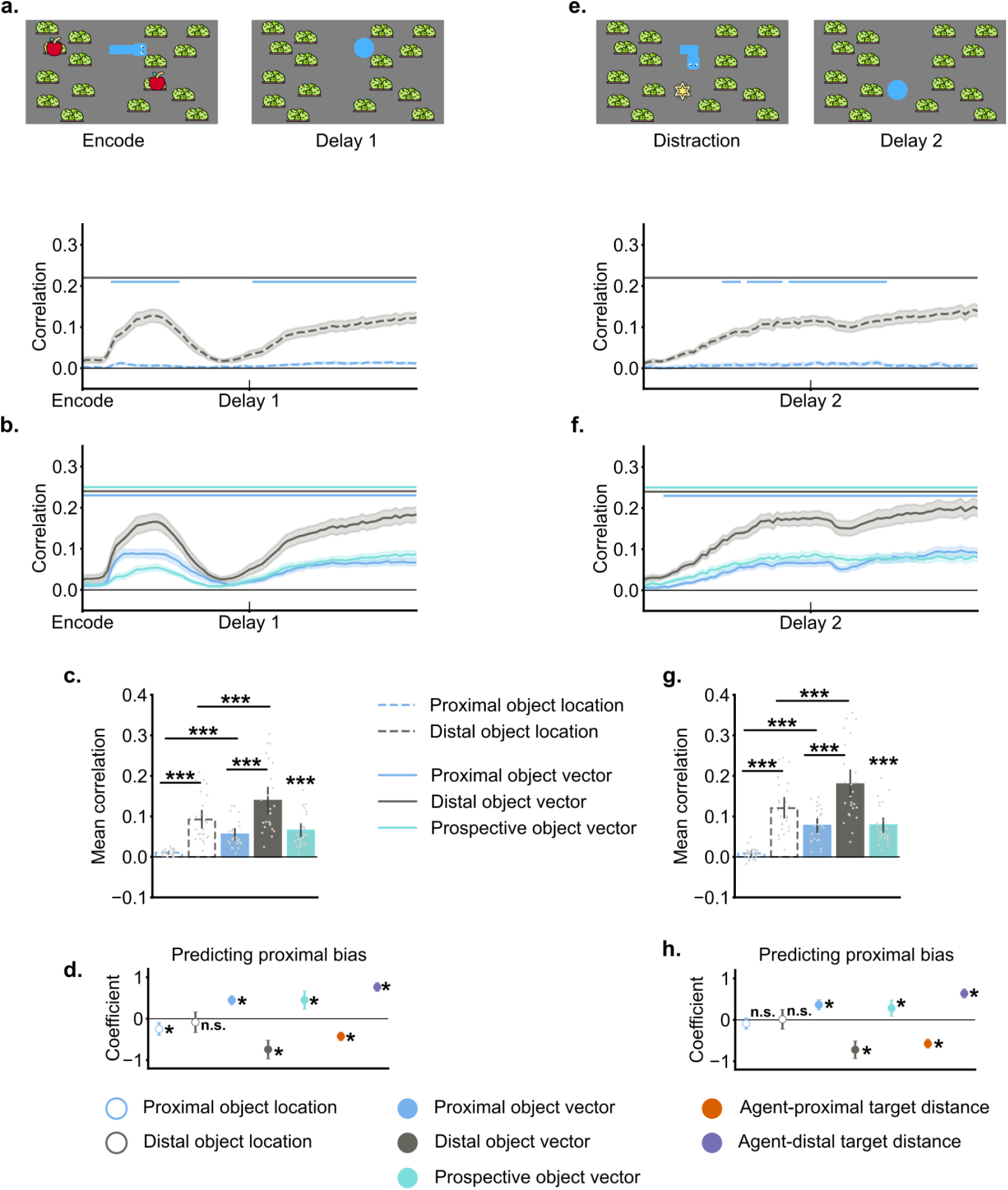
RSA results. a). The time course of object location representations during the encoding and delay 1 periods. b). The time course of object vector representations during the encoding and delay 1 periods. c). Averaged correlations for object locations and object vectors during the delay 1 window. d). GLMM results linking representational strength during the delay 1 window to trial-level proximity bias in no-distraction trials. e). The time course of object location representations during the last 2 s of distraction and delay 2 periods. f). The time course of object vector representations during the last 2 s of distraction and delay 2 periods. g). Averaged correlations for object locations and object vectors during the delay 2 window. h). GLMM results linking representational strength during the delay 2 window to trial-level proximity bias in distraction trials. Error bars represent 95% confidence intervals. * indicates p < .05; *** indicates p < .001. In a,b, e, and f, horizontal colored lines indicate significant evidence for corresponding spatial information. Shaded color represents standard errors.

Instead, it could guide the subsequent movement from the first target to the second. Its representation therefore suggests that participants maintained the spatial relationship between the targets, potentially as part of a multistep action plan.

We next examined which spatial representations contributed to proximity bias, our behavioral index of planning. Trial-level representational strength during the delay 1 window was used to predict whether participants showed a proximity bias on each trial, while controlling for agent-to-target distances (**Fig 3d**). This analysis focused on no-distraction trials because representations maintained during delay 1 were directly relevant to the subsequent memory-guided search in those trials, whereas delay 2 representations were more relevant to search behavior following distraction. As expected, agent-to-target distances strongly modulated proximity bias. As the proximal target became further from the agent, participants were less likely to show a proximity bias, *β* = -0.43, *CI* = [-0.53, -0.33], *BF10* = 10^14^. In contrast, as the distal target became further from the agent, participants were more likely to show a proximity bias, *β* = 0.76, *CI* = [0.66, 0.87], *BF10* = 5. 3 × 10^47^. Among the spatial representations, stronger proximal object vector and prospective object vector coding positively predicted proximity bias, indicating that participants were more likely to select the proximal target first when these representations were stronger (proximal object vector: *β* = 0.45, *CI* = [0.34, 0.56], *BF10* =6. 0 × 10^12^; prospective object vector: *β* = 0.45, *CI* = [0.23, 0.67], *BF10* = 874. 1). In contrast, stronger distal object vector coding predicted a weaker proximity bias, *β* = -0.74, *CI* = [-0.97, -0.52], *BF10* =7. 1 × 10^8^. For object location representations, stronger proximal object location coding negatively predicted proximity bias, *β* = -0.25, *CI* = [-0.40, -0.09], *BF10* = 32.01, whereas distal object location coding was not a reliable predictor, *β* = -0.09, *CI* = [-0.33, 0.16], *BF10* = 0.39.

Overall, object vector coding was more strongly associated with proximity bias than object location coding. Although the distal object vector was represented most strongly during the delay period, stronger distal vector coding predicted a weaker proximity bias. In contrast, stronger proximal and prospective object vector coding predicted a greater proximity bias.

### Object vectors were updated following distraction, reflecting replanning

When the initial action plan was disrupted by the intervening distraction task, participants reinstated the memorized spatial information. This was reflected in a gradual increase in both object vector and object location representations for both targets (**Fig 3e&f**). During the delay 2 window, all spatial representations were reliably represented (*p’s* <= .003) and a repeated-measures ANOVA revealed significance differences in representational strength, *F*(4, 96) = 75.03, *p* < .001, 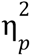 = .48, *BF10* = 3. 89 × 10^25^.

Consistent with the results from delay 1, two main patterns emerged from the pairwise comparisons (**Fig 3g**). First, for both proximal and distal targets, object vector representations were stronger than object location representations [proximal object: *t*(24) = 8.88, *p* < .001, *d* = 2.17, *BF10* = 2. 6 × 10^6^; distal object: *t*(24) = 8.83, *p* < .001, *d* = 0.73, *BF10* = 2. 4 × 10^6^], suggesting a neural bias toward representing target information in relations to the agent position rather than the retrospective object location coordinates. Second, the distal target was represented more strongly than the proximal target, regardless of representational format [object vectors: *t*(24) = 8.31, *p* < .001, *d* = 1.41, *BF10* = 8. 3 × 10^5^; object locations: *t*(24) = 9.05, *p* < .001, *d* = 2.43, *BF10* = 3. 6 × 10^6^], indicating a neural bias toward the distal, future relevant target over the proximal, immediately relevant target. Finally, the prospective object vector was reliably represented, *t*(24) = 8.45, *p* < .001, *d* = 1.69, *BF10* = 2. 2 × 10^6^, indicating that the spatial relation between targets was being tracked. Importantly, because the agent’s position changed during the distraction phase as participants navigated toward the novel star target, the detection of representations of the proximal and distal object vectors indicate that these vectors were updated in memory relative to the agent’s post-distraction position.

We next tested whether these updated object vector representations accounted for the observed proximity bias in distraction trials. Trial-level representational strength during the delay 2 window was used to predict whether participants showed a proximity bias on each trial, while controlling for agent-to-target distances (**Fig 3h**). As expected, agent-to-target distances strongly modulated proximity bias. As the proximal target was located further from the agent, participants were less likely to show a proximity bias for it, *β* = -0.58, *CI* = [-0.68, -0.47], *BF10* = 7. 0 × 10^24^. In contrast, as the distal target was located further from the agent, participants were more likely to show a proximity bias by collecting the proximal target first, *β* = 0.63, *CI* = [0.53, 0.74], *BF10* = 3. 5 × 10^30^. Consistent with the no-distraction trials, stronger representations of the proximal object vector and prospective object vector positively predicted proximity bias, indicating that trials with stronger representations were more likely to end in the optimal collection order (proximal object vector: *β* = 0.37, *CI* = [0.24, 0.49], *BF10* = 3. 6 × 10^6^; prospective object vector: *β* = 0.28, *CI* = [0.09, 0.48], *BF10* = 11. 0). In contrast, stronger distal object vector representations predicted a weaker proximity bias, *β* = -0.72, *CI* = [-0.93, -0.51], *BF10* = 9. 7 × 10^8^. Neither proximal nor distal object location representational strength predicted proximity bias (proximal object location: *β* = -0.09, *CI* = [-0.23, 0.04], *BF10* = 0. 38; distal object location: *β* = 0.01, *CI* = [-0.23, 0.25], *BF10* = 0. 30).

Together, these findings indicate that, following distraction, object vectors were updated relative to the agent’s new position and were represented more strongly than object locations.

Although the distal object vector was represented more strongly than the proximal object vector, stronger distal vector coding predicted a weaker proximity bias, whereas stronger proximal and prospective object vector coding predicted a greater proximity bias.

One possible explanation for the stronger representation of the distal relative to the proximal target is that the distal target posed greater demands on memory fidelity because it had to be maintained for a longer period and across a greater travel distance. Its representation may therefore have been strengthened to support later retrieval rather than immediate action selection. Consistent with this possibility, greater agent-to-target distance, particularly for the distal target, was associated with more visits to non-target locations and greater use of reminders, especially in distraction trials (non-target visits: *β* = 0.19, *CI* = [0.09, 0.28], *BF10* = 219. 87; reminder use: *β* = 0.33, *CI* = [0.16, 0.51], *BF10* = 323. 20). However, we found limited evidence that stronger trial-level representation of the distal target directly reduced memory errors (for full analyses linking representational strength to memory errors, see **SFig 5**). We return to this possibility in the Discussion.

### Memory resources were redistributed post-distraction

Updating action plans after distraction may require memory resources to be redistributed according to the agent’s new position, particularly on trials in which the optimal collection order changed from before to after distraction. On these trials, the item that was initially planned for collection first became the item to be collected second in an updated optimal plan. We hypothesized that changes in memory resource allocation would reflect such action plan updating. To evaluate this idea, we compared item-level representational strengths before and after distraction. Trials were divided into a *stay* condition, in which the optimal collection order remained unchanged, and a *switch* condition, in which the optimal order changed following distraction. We conducted a repeated-measures ANOVA with factors of time (pre-distraction vs. post-distraction), trial type (stay vs. switch), and spatial representation (initially proximal object location, initially proximal object vector, initially distal object location, and initially distal object vector). This analysis revealed a significant three-way interaction, *F*(3, 72) = 77.76, *p* < .001, 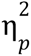 = .76, *BF10* = 7. 5 × 10^40^. Paired t-tests were then used to assess changes in representational strength from before to after distraction. On switch trials (**Fig 4c&d**), the initially proximal item became the distal item after distraction, and its representational strength increased for both object location and object vector codes [proximal object location: *t*(24) = 8.12, *p* < .001, *d* = 1.93, *BF10* = 5. 7 × 10^5^; proximal object vector: *t*(24) = 8.66, *p* < .001, *d* = 2.03, *BF10* = 7. 6 × 10^9^]. In contrast, the initially distal item became the proximal item after distraction, and its representational strength decreased for both object location and object vector codes [distal object location: *t*(24) = -8.02, *p* < .001, *d* = 1.82, *BF10* = 4. 6 × 10^5^; distal object vector: *t*(24) = -7.05, *p* < .001, *d* = 1.39, *BF10* = 5. 9 × 10^4^]. On stay trials (**Fig 4a&b**), there was no evidence of a comparable redistribution of memory resources between the two items. The initially distal item remained more strongly represented than the initially proximal item both before and after distraction. Nevertheless, overall representational strength increased from delay 1 to delay 2 for the proximal target [location:*t*(24) = -0.28, *p* = 1.00, *BF10* = 0. 22; vector: *t*(24) = 7.30, *p* < .001, *d* = 1.11, *BF10* = 1. 0 × 10^5^], and distal target [location: *t*(24) = 7.63, *p* < .001, *d* = 1.26, *BF10* = 5. 8 × 10 ; vector: *t*(24) = 8.90, *p* < .001, *d* = 1.06, *BF10* = 5. 9 × 10^8^]. Together, these findings suggest that memory representations were selectively reweighted to support optimal action replanning following distraction.

**Fig. 4.**
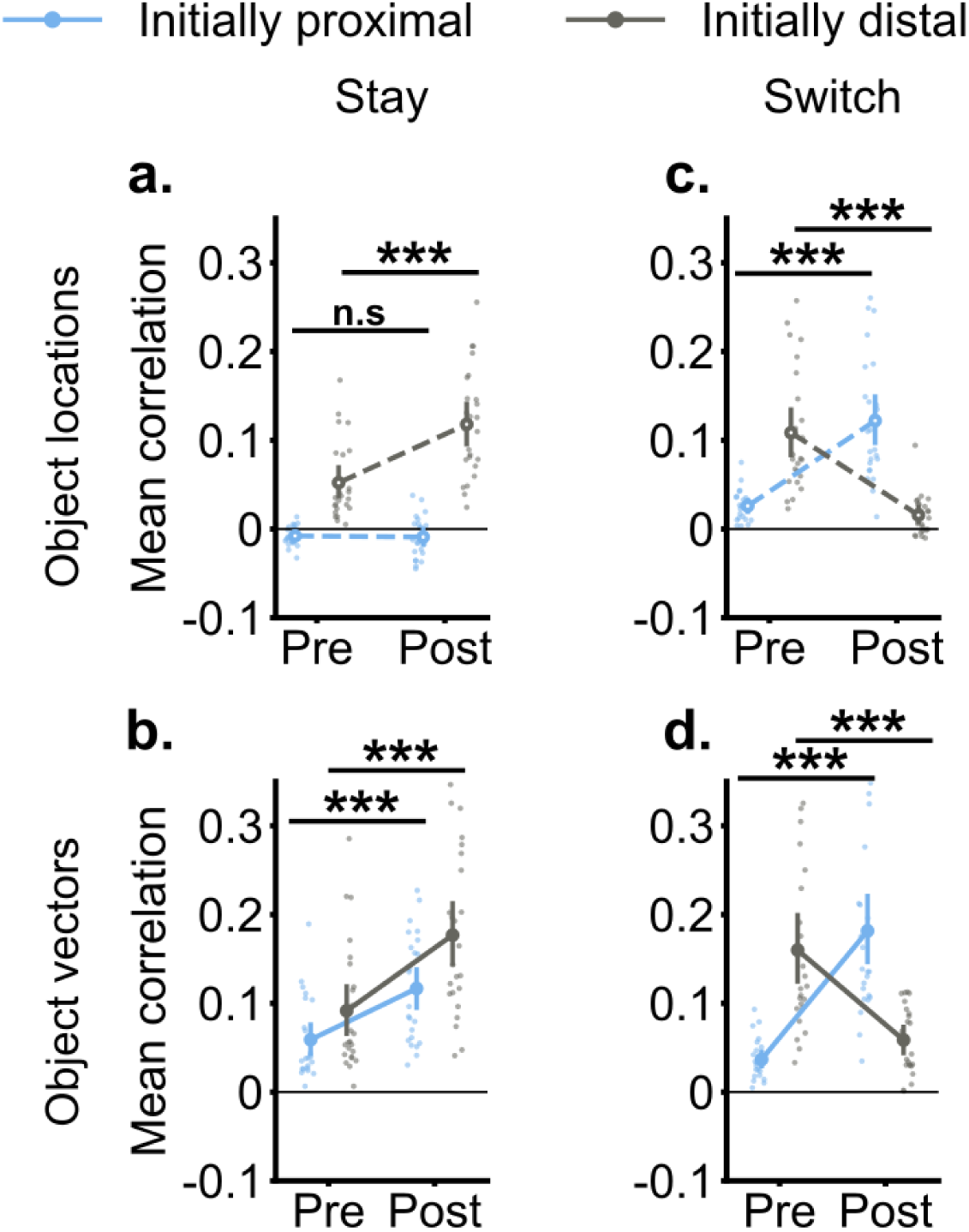
Representational strengths before and after distraction. In stay trials, pre and post distraction representational strength for a) object locations and b) object vectors. In switch trials, pre and post distraction representational strength for a) object locations and b) object vectors. n.s. indicates p > .05; *** indicates p < .001. Error bars represent 95% confidence intervals.

### External attention was biased toward the proximal (immediate) target despite stronger neural representations of the distal (future) target

The finding that distal targets were represented more strongly neurally than proximal targets is intriguing because it contrasts with the observed behavioral proximity bias. One possible mechanism that may help reconcile this paradox is the allocation of external attention through eye movements. We therefore tested whether gaze during the delay periods was biased toward either target location, even though participants were explicitly instructed to maintain fixation at the agent’s position. During encoding and delay 1 periods (**Fig 5a**), fixations were more likely to fall near the proximal than the distal target, after controlling for differences in agent-to-target distance (see Methods). In delay 1, the mean proportion of fixations on the proximal target was significantly greater than that on the distal target (**Fig 5b**), *t*(23) = 3.00, *p* = .006, *d* = 0.73, *BF10* = 7.06. A similar pattern emerged during the distraction and delay 2 periods (**Fig 5c**). Fixations were again more likely to fall near the proximal target than the distal target. During delay 2, the mean proportion of fixations on the proximal target was significantly greater than that on the distal target (**Fig 5d**), *t*(23) = 3.95, *p* <.001, *d* = 1.28, *BF10* = 52.57.

**Fig. 5.**
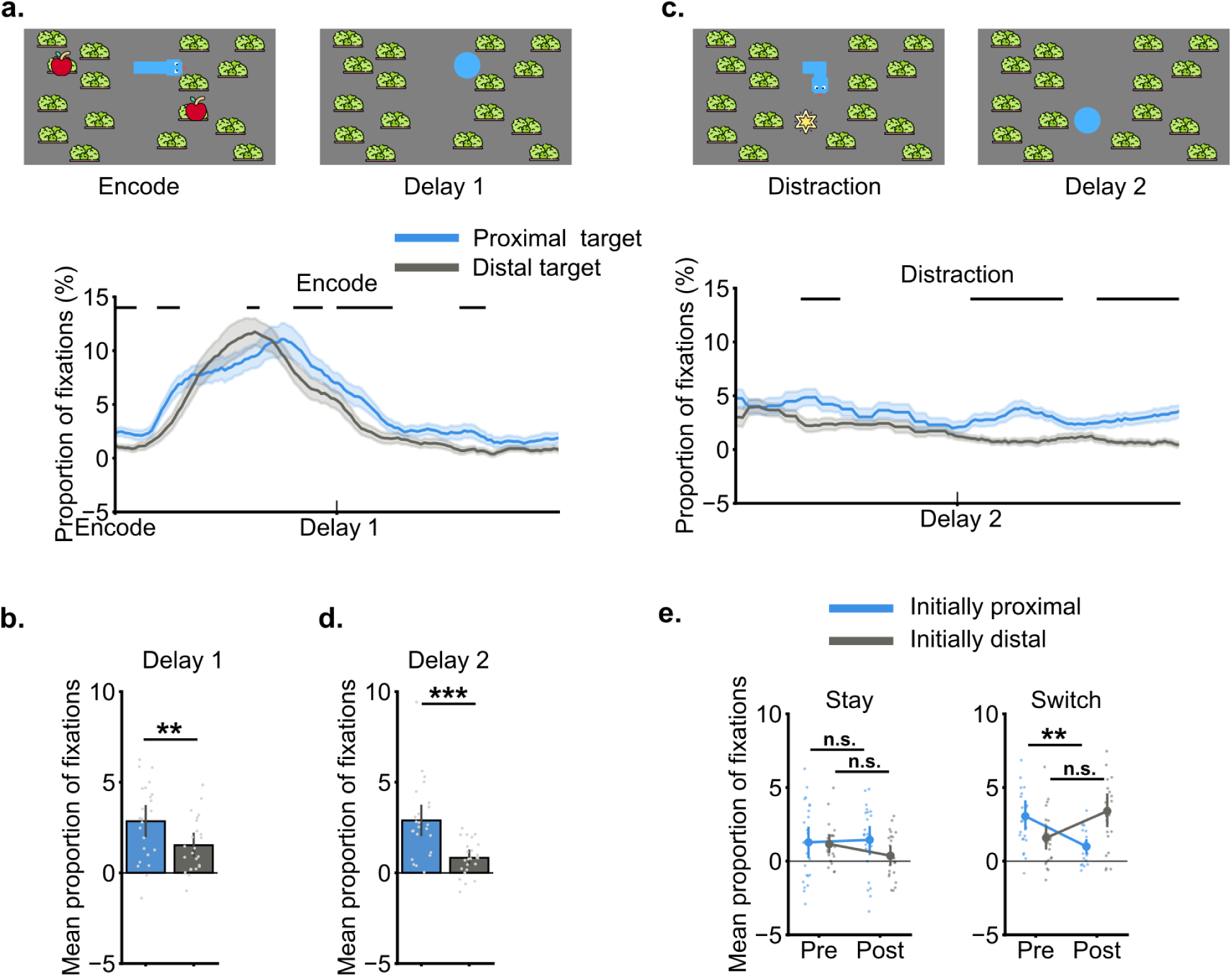
Fixation results. a). Fixations on object locations as a function of time during encoding and the first delay window. b). During the delay following encoding, more fixations landed on proximal objects than on distal objects. c). Fixations on object locations as a function of time during the last two seconds of the distraction phase and the second delay window. d). During the delay following distractions, more fixations landed on updated proximal objects than on updated distal objects. e). Changes in proportions of fixations on items pre-distraction and post-distraction in stay and switch trials. Error bars indicate 95% confidence intervals. n.s indicates p > .05; ** indicates p < .01. *** indicates p < .001. In a and c, horizontal colored lines indicate significant differences between the proximal and distal targets. The shaded area represents standard errors.

We next compared fixation patterns between stay and switch trials. This analysis tested whether replanning was accompanied by corresponding shifts in fixation bias. A repeated-measures ANOVA with factors of time (pre-distraction vs. post-distraction), trial type (stay vs. switch), and item (initially proximal vs. initially distal) revealed a significant three way interaction, *F*(1, 23) = 13.11, *p* = .001, 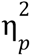 = .36, *BF10* = 341.56. Follow-up ANOVAs revealed a significant interaction between item and time for switch trials, *F*(1, 23) = 14.07, *p* = .001, 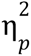 = .18, *BF10* = 3. 3 × 10^5^, but not stay trials, *F*(1, 23) = 1.69, *p* = .418, 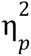 = .01, *BF10* = 0.15. On switch trials (**Fig 5e**), the initially proximal item became the distal item after distraction, and the proportion of fixations landing on it decreased [*t*(23) = -3.61, *p* = .006, *d* = 1.08, *BF10* = 24.82]. In contrast, the initially distal item became the proximal item after distraction, and there is weak evidence that the proportion of fixations landing on it may have increased, *t*(23) = 2.53, *p* = .073, *d* = 0.79, *BF10* = 2.86. On stay trials, no significance difference between pre-distraction and post-distraction was found [initially proximal object: *t*(23) = 0.26, *p* = .998, *BF10* = 0.22; initially distal object: *t*(23) = 2.10, *p* = .173, *BF10* = 1.38]. These results were robust to variability in the empirical sampling rate (**SFig 6**).

### Dynamic reconfiguration of action plans

So far we have analyzed neural and behavioral indices of replanning by comparing pre-distraction and post-distraction periods. The early emergence of updated object vectors suggests that replanning happened during the distraction period rather than at the end of it. To evaluate how replanning unfolds over time, we analyzed eye movement trajectories.The updated optimal action plan in our task was to move from the updated agent position first toward the proximal target and then to the distal one. We therefore defined forward and backward gaze “sweeps” for both optimal and suboptimal plans relative to the updated agent position at the end of the distraction period (i.e., at the position of the star target). We also included gaze patterns associated with the independent encoding of the two targets (**Fig 6a&b**). A repeated-measures ANOVA revealed a main effect of gaze patterns, *F*(5, 115) = 28.23, *p* < .001, 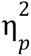 = .57, *BF10* = 3. 9 × 10^25^. Planned *t*-tests compared the occurrence rates of paired gaze patterns (**Fig 6c**). Backward sweeps aligned with optimal plans occurred more frequently than those aligned with suboptimal plans, *t*(23) = 3.53, *p* = .002, *d* = 0.71, *BF10* = 20.98). This pattern of eye gazes suggests that participants actively reconfigured efficient action plans while completing the distraction task. No significant difference between optimal and suboptimal forward sweeps was found, *t*(23) = -1.23, *p* = .232, *BF10* = 0.42. Gaze patterns reflecting the independent encoding of individual items occurred more frequently for proximal than for distal targets, indicating that proximal targets were more likely to be encoded relative to the agent’s position, *t*(23) = 6.30, *p* < .001, *d* = 1.49, *BF10* = 9520.97. We tested whether the rate of each gaze pattern predicted optimal behavior on a given trial, but found no reliable associations (**Sfig 7a**).

**Fig. 6.**
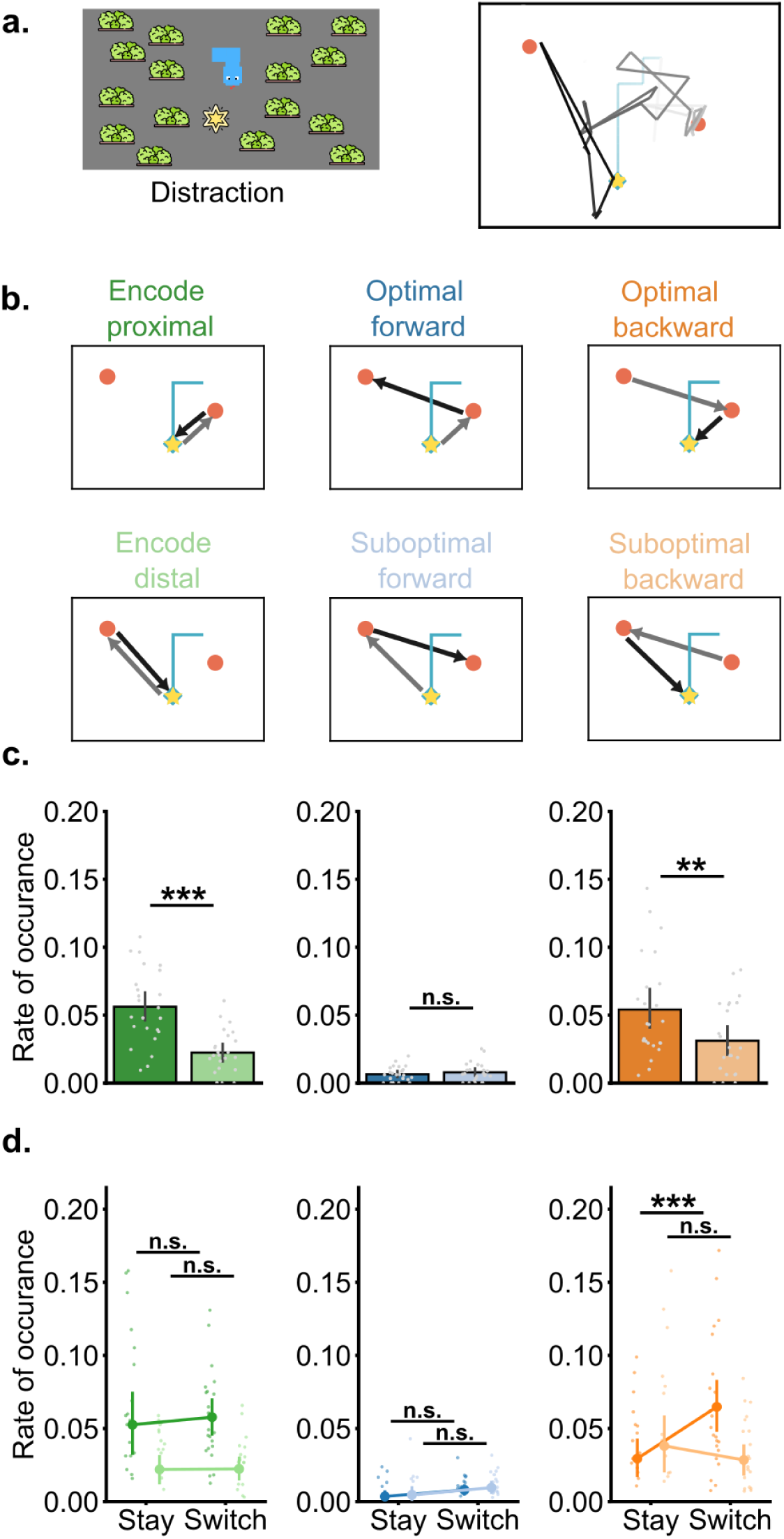
Gaze trajectories during the distraction phase. a). Example gaze trajectories, with lighter colors indicating earlier time points and darker colors indicating later time points. b). Illustrations of gaze patterns in an example target display. c). Rate of occurrence for each gaze pattern. d). Rate of occurrence for each gaze pattern for stay and switch trials. n.s indicates p > .05; ** indicates p < .01; *** indicates p < .001. Error bars represent 95% confidence intervals. [dist_combined.mp4](https://drive.google.com/file/d/1CbB4NoD0CIVIIj_qN6hdkjGJhl6TR9jb/view?usp=sharing) **Video. 3.** Illustrations of gaze trajectories during the distraction and delay 2 periods from five example trials.

If backward sweeps reflect online replanning during the distraction phase, they should occur more frequently when replanning is required – on switch trials, in which the optimal collection order changed following distraction. We therefore compared the occurrence rates of each gaze pattern between stay and switch trials. A repeated-measures ANOVA with variables of trial type (stay vs. switch) and gaze pattern types revealed a significant interaction, *F*(5, 115) = 4.49, *p* < .001, 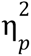= .16, *BF10* = 14.15. Follow-up comparisons showed that only backward sweeps occurred more frequently on switch trials than on stay trials, *t*(24) = 5.21, *p* < .001, *d* = 0.94, *BF10* = 857.95. No other gaze patterns differed significantly between trial types (*ps* > 0.3). We conducted control analyses and confirmed that these findings were robust to variability in the empirical sampling rate (**Sfig 7b&c**).

## Discussion

Classic working memory research has primarily emphasized its retrospective role in maintaining and manipulating previously encountered information (Baddeley 2003; Luck and Vogel 1997, 2013). More recently, growing attention has been directed toward its prospective functions, particularly how working memory interacts with planning and decision-making to guide goal-directed behavior in dynamic and naturalistic environments (Boettcher et al., 2021; Draschkow et al. 2021; van Ede et al. 2019; van Ede and Nobre 2023). Building on this emerging perspective, the present study addressed two foundational questions: What representations support the formation of spatial action plans in working memory, and how do these representations enable flexible adaptation to changes in the environment?

Using a working memory paradigm inspired by the classic arcade game *Snake*, we found that participants configured efficient action plans for collecting target apples, typically prioritizing the target proximal to the agent. When an intervening distraction changed the agent’s position, participants flexibly updated their plans, shifting priority toward targets that were proximal to the new position rather than rigidly adhering to the original collection order. Eye movement analyses showed that participants engaged in gaze trajectories consistent with action planning during encoding. RSA of the EEG data further revealed that object vector representations (reflecting the coordinates of each apple target relative to the snake agent) were maintained more strongly than absolute object location representations during the delay, which reliably predicted behavioral proximity biases. These representations included vectors linking the agent to individual targets as well as a prospective vector capturing the spatial relationship between the two targets. Following distractions that disrupted the original plan, replanning occurred by updating agent-centered object vectors relative to the agent’s new position. Gaze trajectories revealed that the replanning process unfolded dynamically during the distraction phase. Together, these findings suggest that working memory maintained spatial information in a dynamic, action-relevant format that was actively reconfigured when task demands changed. In the sections that follow, we organize the findings according to the task phase in which they emerged, providing an integrated account of the processes and representations that supported planning and flexible replanning in this dynamic task paradigm.

During encoding, eye-movement trajectories suggested that planning was closely linked to the sequential sampling of spatial information. In spatial navigation studies, both forward sweeps, extending from the current position toward a goal, and backward sweeps, tracing a route from the goal back toward the current position, have been associated with effective route planning (Zhu et al. 2022). Similar principles have been observed in non-spatial domains, where plans can be constructed either forward from the current state or backward from a desired outcome (Ambrose, Pfeiffer, and Foster 2016; Liu et al. 2019; Park, Lu, and Hedgcock 2017).

Consistent with this broader literature, participants in our task exhibited systematic gaze trajectories during encoding. The proximal target was more likely to be sequentially linked with the agent’s position than the distal target. Moreover, gaze was more likely to follow the optimal sequence from the agent to the proximal target and then to the distal target than the corresponding suboptimal sequence in reverse order. These patterns suggest that participants were already configuring efficient multistep plans during encoding.

During the delay window following encoding, we evaluated how the configured plan was maintained. The stronger representation of object vectors relative to object locations indicates a neural bias toward coding target locations in relation to the agent. The concepts of object location and object vector codes are grounded in the spatial navigation literature. In rodents, object vector representations formed in hippocampal theta sweeps have links to goal-directed planning during spatial navigation (Ormond and O’Keefe 2022; Tang et al. 2026; Yu et al. 2026). Although our paradigm differs from natural navigation because the agent and targets were presented on a two-dimensional display, these concepts provide a useful framework for distinguishing absolute spatial codes from relational, agent-centered codes. Object vectors contain information that is directly relevant for action in our task, including the direction and distance from the agent to a target. Their stronger representation may therefore reflect a transformation of remembered sensory information into a prospective format better suited for guiding future behavior. Previous studies have often dissociated stimulus features from motor requirements to demonstrate prospective action coding in working memory (Boettcher et al., 2021; van Ede et al. 2019; Ehrlich and Murray 2022; Henderson et al. 2022; Nasrawi et al. 2023). Our findings extend this work by showing that prospective coding can also support more abstract, multistep plans involving relations among the agent and multiple future targets.

One alternative interpretation is that the early emergence of object vector coding during encoding may reflect the natural relational encoding of spatial layouts rather than prospective planning per se. Several additional observations nevertheless support a functional role for these representations in action guidance. First, trial-level object vector strength was related to participants’ subsequent target collection order. Stronger proximal and prospective object vector representations were associated with a stronger proximity bias, i.e., a greater likelihood of selecting the proximal target first, whereas stronger distal object vector representations predicted a weaker proximity bias. Second, when a distraction task interrupted the original plan, object vectors were recomputed relative to the agent’s new position in memory, even though the absolute target locations remained unchanged. Thus, object vector coding was not merely a static property of the visual display. Instead, it was dynamically updated in a manner that reflected the spatial information relevant to the participant’s subsequent actions.

A notable finding was that the distal target had stronger neural representation than the proximal target, despite participants’ behavioral tendency to collect the proximal target first. Eye movements showed the opposite pattern: gaze was preferentially directed toward the proximal target during both the pre- and post-distraction delay periods. This dissociation suggests that internal working memory representations and external attention may make simultaneous, complementary contributions to multistep planning (Ballard et al. 1995; Draschkow et al. 2021; Hayhoe and Ballard 2005; Keshava et al. 2024). Prior navigation studies have shown that visual attention is often directed toward immediately relevant transitions rather than final goal locations, consistent with a close coupling between external attention and the next action (Keshava et al. 2024; Zhu et al. 2022). In our task, gaze may similarly have prioritized the proximal target because it was most relevant to the immediately upcoming movement.

The stronger neural representation of the distal target may reflect a different functional demand. One possibility is that the distal target served as the ultimate goal of the multistep plan and therefore required stronger maintenance while attention was directed toward the subgoal of collecting the proximal target. A related possibility is that the distal target imposed greater demands on memory fidelity because it had to be retained for longer and across a greater travel distance. Consistent with this account, greater agent-to-target distance, particularly for the distal target, was associated with more memory errors and more reliance on reminders. However, we found limited evidence that stronger trial-level representation of the distal target directly reduced such errors. Future work could distinguish whether this pattern reflects a tendency of stronger prospective goal coding, compensatory allocation of memory resources, or some other aspect of multistep planning.

When initial plans were disrupted by distraction, object vectors were recomputed relative to the agent’s new position in the post-distraction delay window. The strength of these updated representations predicted proximity bias on distraction trials, further supporting their role in guiding future actions. In addition to updating the representational content, memory resources also appeared to have been redistributed to accommodate the revised action plan. Distal targets were generally represented more strongly than proximal targets, and this reallocation was especially evident on switch trials, in which the optimal collection order changed after distraction. On these trials, the representational strength of a given item shifted according to its new role in the plan: items that became newly proximal were represented less strongly, whereas items that became newly distal were represented more strongly. Prior studies examining post-distraction neural representations have typically focused on their robustness or recovery (Bettencourt and Xu 2016; Hallenbeck et al. 2021; Mallett and Lewis-Peacock 2019; Rademaker et al. 2019). Our findings extend this work by revealing a broader pattern of representational change. Rather than being merely preserved or reinstated, internal representations were actively transformed and recomputed to accommodate changes in the environment, while memory resources were redistributed in accordance with the demands of the updated plan.

In addition to shifts in neural representations following distraction, gaze was biased toward the newly proximal target, directly contrasting with the neural bias toward the distal target. This reallocation of gaze as the action plan was updated was observed most clearly in the comparison between stay and switch trials. On switch trials, in which the optimal collection order changed following distraction, gaze shifted away from the target that had initially been proximal and toward the distal target, which had become newly proximal. Together, the redistribution of memory resources, reflected in changes in neural representational strength, and the reallocation of attentional resources, reflected in changes in gaze patterns, suggest complementary roles for working memory and attention in multistep planning. Although the interaction between attention and memory has long been a central topic across multiple fields, how these systems work together to support behavior in dynamic, naturalistic environments remains an important open question. Our findings highlight a potentially complementary division of labor between these processes that should be examined more directly in future research.

Finally, we turn to the pivotal question of how replanning unfolded. Gaze trajectory analyses indicated that participants actively reconfigured their action plans *during* the distraction period, relying primarily on backward sweeps that revisited the memorized target locations. This pattern revealed two key features of the replanning process. First, replanning was active.

Although participants were never instructed to collect the targets in a particular order, they did not rigidly adhere to their initial plans. Instead, they actively resampled memorized spatial information during the distraction period and flexibly incorporated changes in the environment into updated action plans. Second, replanning unfolded dynamically while the distraction task was still ongoing. Rather than waiting until the distraction had ended, participants proactively revisited the memorized target locations while continuing to pursue the orthogonal goal of the distraction task. This pattern contrasts with accounts in which recovery of working memory representations is treated primarily as a post-distraction process (Hallenbeck et al. 2021; Kozachkov et al. 2022; Mallett and Lewis-Peacock 2019). In naturalistic environments, multiple goals often compete for control of behavior. In our task, the sudden subgoal of reaching the distractor target was deliberately designed to compete with the prospective goal of collecting the target apples. Participants did not appear to treat these goals as strictly sequential and independent. Instead, they dynamically integrated them. Eye movements showed that participants sampled the two memorized target locations before shifting gaze toward the agent’s future position, which coincided with the distractor. This strategic pattern suggests that planning for the future goal was embedded within pursuit of the current one. An important question for future research is whether backward sweeps would still emerge if the agent’s future position were spatially dissociated from the distractor. Such a manipulation would help disentangle replanning for the future goal from processes directly related to completing the current distraction task.

Our findings align with the emerging view that working memory is fundamentally prospective, serving not only as a store of past sensory information but also as a system for preparing and guiding future actions (Boettcher et al., 2021; van Ede et al. 2019; Ehrlich and Murray 2022; Henderson et al. 2022; Nasrawi et al. 2023; Rainer et al. 1999). Previous research has largely examined this perspective in settings in which distinct sensory stimuli were mapped onto specific motor responses, allowing retrospective sensory codes and prospective motor codes to be tracked separately (Boettcher et al., 2021; van Ede et al. 2019; Henderson et al. 2022; Nasrawi et al. 2023). These studies have shown that, in addition to preserving sensory information, working memory also carries prospective response codes, indicating that action preparation can begin during or shortly after encoding. Our study extends this framework beyond stimulus response mappings by examining how remembered sensory information is transformed into multistep action plans. Importantly, we used a dynamic and relatively unconstrained environment in which plans could develop naturally and had to be revised as task demands changed. This approach shows that working memory not only supports the formation and maintenance of prospective plans, but also flexibly reconfigures those plans when environmental perturbations occur, highlighting its role as a dynamic control system for adaptive behavior.

In conclusion, participants configured efficient multistep plans for collecting targets and flexibly updated those plans when distraction displaced the agent. Rather than adhering to their initial collection order, they prioritized targets that became newly accessible from the agent’s updated position. EEG analyses revealed a representational bias toward dynamic object vectors over static object locations, including prospective vectors that captured spatial relationships between successive targets. Following distraction, agent-centered vectors were updated relative to the new agent position, and item-level representational strength was redistributed when the optimal target order changed. Eye-movement analyses further showed that gaze was biased toward the immediately relevant target, whereas memory representations were biased toward the future-relevant target, suggesting simultaneous, complementary functional roles for attention and memory in spatial planning. Structured gaze sweeps accompanied both initial planning during encoding and dynamic replanning during the distraction phase. Together, these findings highlight working memory as a dynamic, prospectively oriented system that reorganizes remembered information to guide adaptive behavior in changing environments.

---

**SFig. 1.**
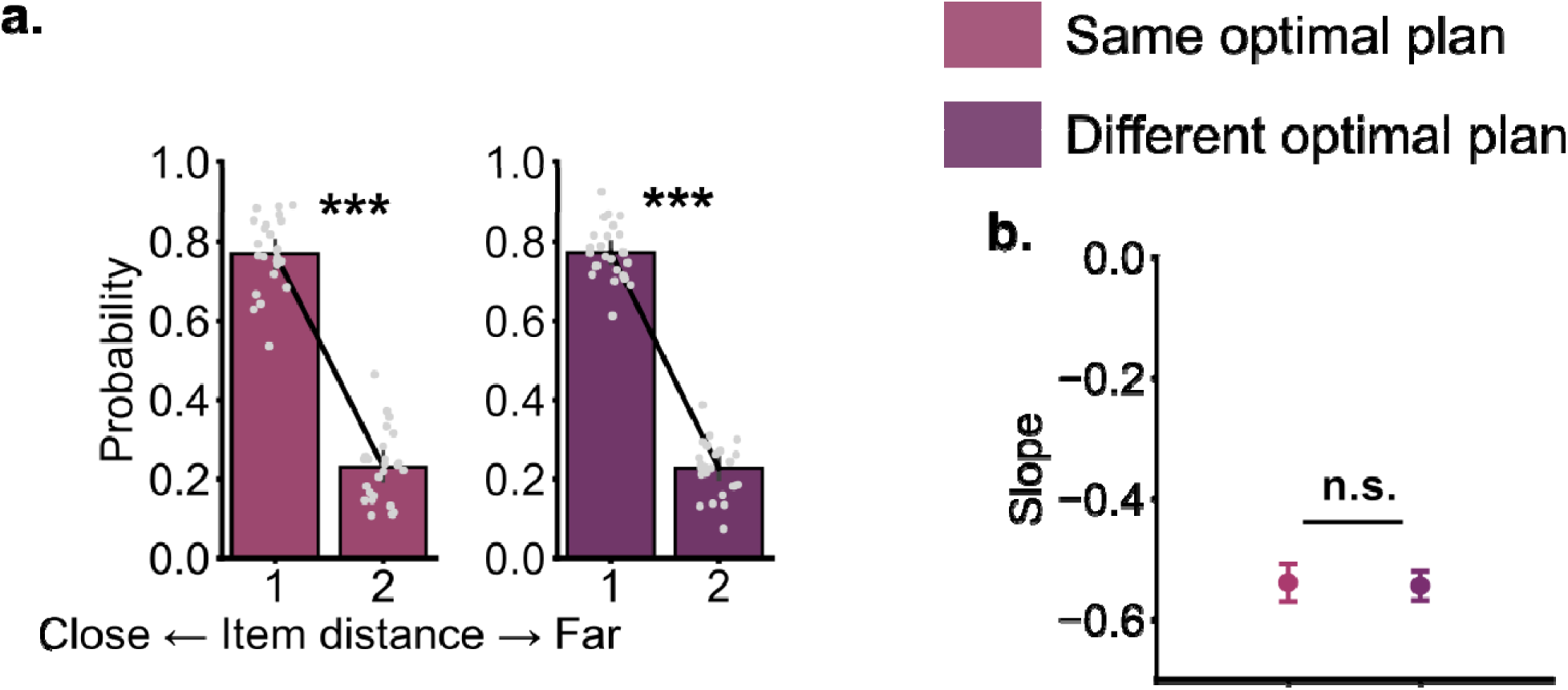
Behavioral proximity bias in distraction trials. a). Probability of an item being collected as the first target as a function of its relative distance to the agent’s position in trials where the optimal plan should have been updated following distraction or trials where the optimal plan remained the same. f). Slopes of linear models fitted to the probability of an item being collected first as a function of distance. Error bars represent 95% confidence intervals. n.s. indicates p > .05; *** indicates p < .001.

## Additional analyses of gaze trajectories during the encoding window

To examine the functional relevance of these gaze patterns, we used the rate of gaze patterns in a trial to predict whether that trial ended with showing optimal behavior in no-distraction trials (**SFig 2a**). Gaze patterns associated with independently encoding of the proximal target positively predicted proximal bias (β = 0.12, *CI* = [0.01, 0.24], *BF10* = 1.59), whereas gaze patterns associated with independently encoding of the distal target negatively predicted proximal bias (β = -0.11, *CI* = [- 0.20, -0.02], *BF10* = 1.83). Forward and backward sweep patterns did not reliably predict trial level behaviors (optimal forward sweep, β = 0.10, *CI* = [-0.00, 0.21], *BF10* = 0.73; suboptimal forward sweep, β = -0.08, *CI* = [-0.18, 0.01], *BF10* = 0.66; optimal backward sweep, β = 0.00, *CI* = [-0.10, 0.11], *BF10* = 0.14; suboptimal backward sweep, β = 0.00, *CI* = [-0.10, 0.11], *BF10* = 0.14).

We conducted control analyses to assess whether gaze pattern findings were robust to variability in the empirical sampling rate (**SFig 2b**). Specifically, we repeated the analyses after resampling each participant’s eye-tracking data to their lowest observed empirical sampling rate. A repeated-measures ANOVA revealed a significant main effect of gaze pattern, *F*(5, 115) = 8.36, *p* < .001, = .47. At this reduced sampling rate, we replicated the main finding that forward sweeps that were aligned with optimal plans occurred more frequently than those aligned with suboptimal plans, but not for backward sweeps (optimal vs. suboptimal forward sweeps: *t*(23) = 2.25, *p* = .034, *d* = 0.32, *BF10* = 1.76; optimal vs. suboptimal backward sweeps: *t*(23) = 1.80, *p* = .085, *d* = 0.27, *BF10* = 0.86). In addition, gaze patterns consistent with the independent encoding of the proximal target occurred more frequently than those associated with the distal target, *t*(23) = 2.29, *p* = .032, *d* = 0.48, *BF10* = 1.87.

**SFig. 2.**
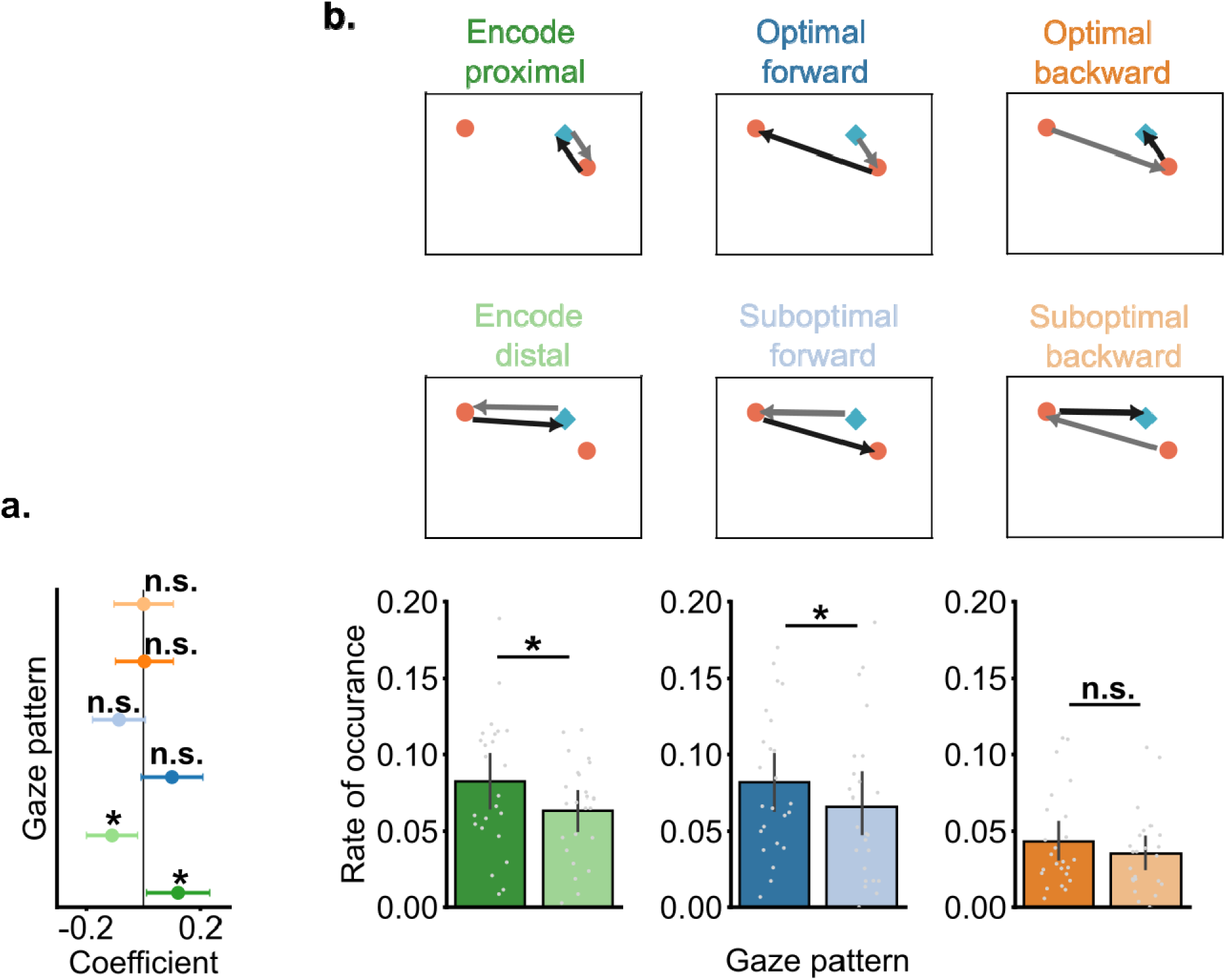
Additional analyses of gaze trajectories during the encoding window. a). Trial-level GLMM relating the rate of each gaze pattern to whether participants subsequently selected the optimal collection order in no-distraction trials. b). Rate of each gaze pattern after resampling the eye-tracking data to a reduced sampling rate. Error bars represent 95% confidence intervals. n.s indicates p > .05; * indicates p < .05.

**SFig. 3.**
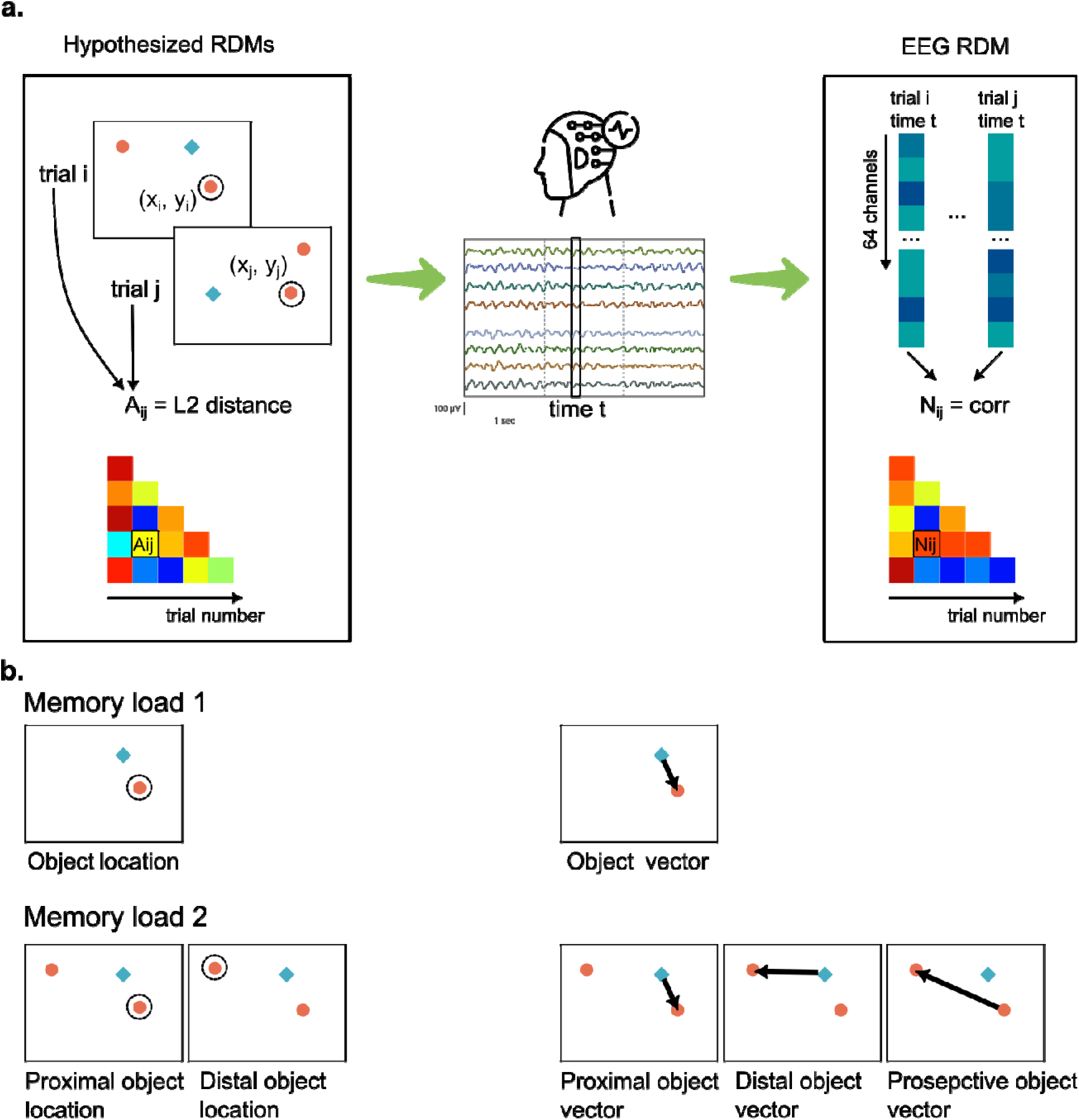
RSA approach and spatial features. a). Schematic of hypothesized representational dissimilarity matrices (RDMs) and corresponding neural RDMs. b). Spatial features included in the RSA analyses, including object locations and object vectors, for memory load 1 and load 2 trials.

## Object location and object vector coding in memory load 1 trials

We first examined if object location and object vector were represented in EEG patterns in memory load 1 trials. For the encoding window, we constructed model RDMs for the target position, and object vector. Both object location and object vector representations emerged during the encoding and delay 1 window (**SFig 4a**). During the delay window, there were reliable object location and object vector representations, and object vector representations were stronger than object location [**SFig 4b**, object vector: *t*(24) = 8.27, *p* < .001, *d* = 1.65, *BF10* = 1.5 × 10^6^; object location: *t*(24) = 6.10, *p* < .001, *d* = 1.22, *BF10* = 1.5 × 10^4^; object vector vs object location: *t*(24) = 8.80, *p* < .001, *d* = 1.26, *BF10* = 2.2 × 10^6^]. We next tested whether mean representational strength predicted trial-level memory errors in no-distraction trials. For non-target visits, greater agent-object distance predicted more non-target visits, β = 0.30, *CI* = [0.00, 0.59], *BF10* = 2.52. Although object location strength did not predict non-target visits, β = 0.42, *CI* = [-0.13, 0.97], *BF10* = 2.16, object vector strength predicted fewer non-target visits, β = -0.81, *CI* = [-1.25, -0.29], *BF10* = 76.05. Neither representational strength nor agent-target distance reliably predicted reminder use in no-distraction trials (**SFig 4c**). The effects were as follows: agent-object distance, β = 0.23, *CI* = [-1.93, 2.26], *BF10* = 2.72; object location, β = 0.62, *CI* = [-2.37, 4.18], *BF10* = 4.30; object vector, β = -1.71, *CI* = [-5.31, 1.29], *BF10* = 6.06.

For the distraction and post-distraction periods, we constructed model RDMs corresponding to target position, the updated object vector from the updated agent position to the target, and the initial object vector from the initial agent position to the target. These models allowed us to test whether object location coding emerged after distraction and whether the object vector representation was updated to reflect the agent’s new position. Reliable object location and updated object vector representations emerged and sustained during the late distraction phase and the delay 2 phase (**SFig 4d**). During the post-distraction delay window, a significant difference in representational strength among object locations, updated object vectors, and initial object vectors was found, *F*(2, 48) = 87.19, *p* < .001, 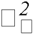 = .57. Updated object vector coding was stronger than for either object locations or initial object vectors [**SFig 4e**, updated object vector: *t*(24) = 9.92, *p* < .001, *d* = 1.98, *BF10* = 3.7×10^7^; object location: *t*(24) = 7.68, *p* < .001, *BF10* = 4.5×10^5^; initial object vector: *t*(24) = 2.64, *p* = .007, *d* = 0.53, *BF10* = 7.04; updated object vector vs object location: *t*(24) = 8.98, *p* < .001, *d* = 1.36, *BF10* = 3.2×10^6^; updated object vector vs initial object vector: *t*(24) = 9.85, *p* < .001, *d* = 2.57, *BF10* = 1.7×10^7^].

Finally, we tested whether post-distraction representational strength predicted trial-level memory errors, including reminder use and non-target visits. Greater agent-target distance predicted more non-target visit errors (β = 0.33, *CI* = [0.14, 0.51], *BF10* = 104.63), whereas stronger updated object vector representations predicted fewer non-target visits (**SFig 4f**, β = - 0.36, *CI* = [-0.66, -0.07], *BF10* = 6.14). No other reliable associations were observed. For reminder use, the effects were: agent–target distance, β = 0.45, *CI* = [-0.07, 0.97], *BF10* = 2.99; object location, β = -0.02, *CI* = [-1.08, 1.08], *BF10* = 1.54; updated object vector, β = -0.33, *CI* = [-1.29, 0.64], *BF10* = 1.61; initial object vector, β = 0.21, *CI* = [-0.40, 0.86], *BF10* = 1.00]. For non-target visits, the effects were: object location, β = -0.15, *CI* = [-0.47, 0.17], *BF10* = 0.61; initial object vector, β = 0.11, *CI* = [-0.09, 0.32], *BF10* = 0.45).

**SFig. 4.**
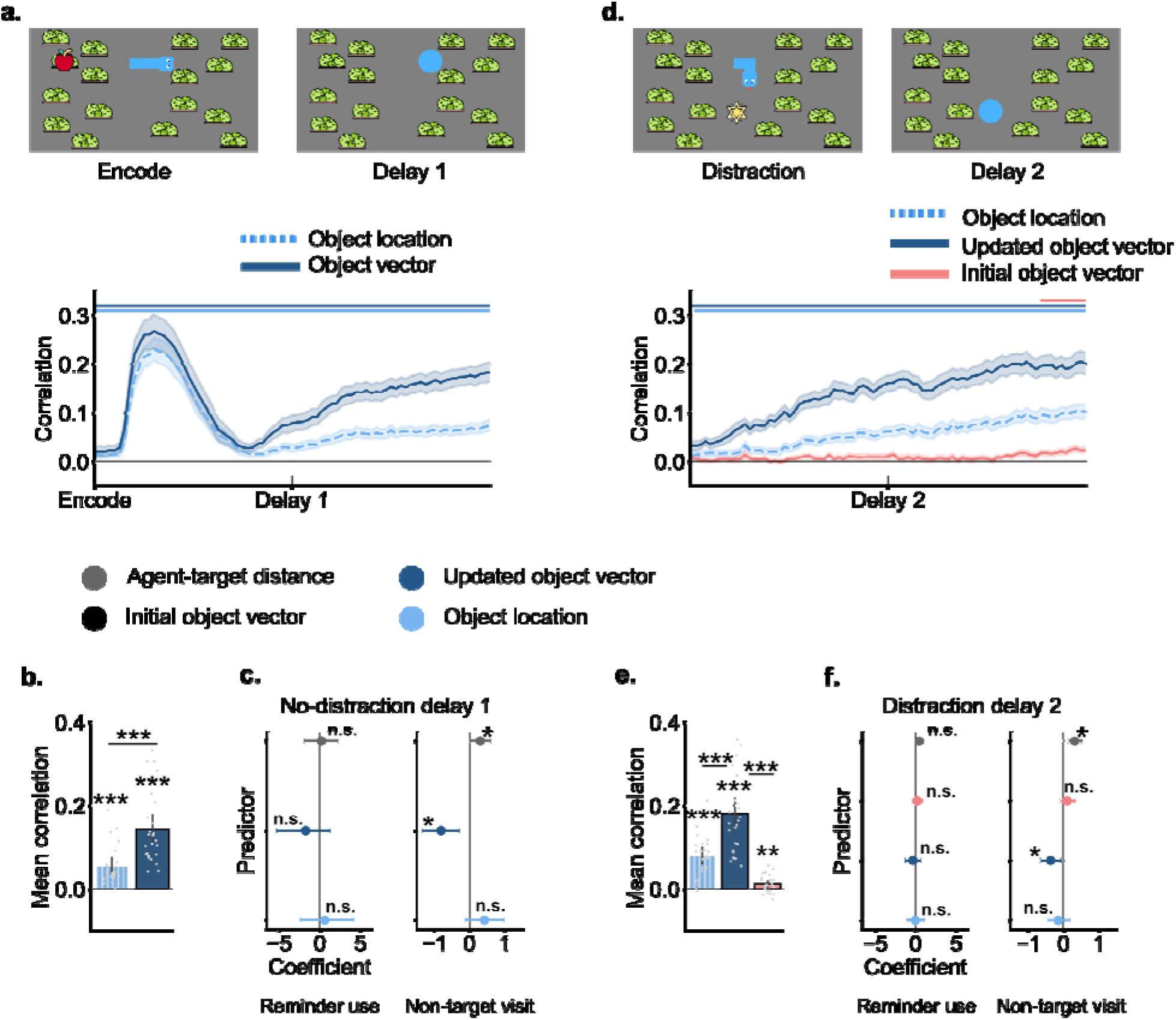
Object location and object vector coding in memory load 1 trials. a). Time courses of object location and object vector strength during encoding and delay 1. b). Mean representational strength during delay 1. c). Trial-level GLMM testing whether representational strength of object location, object vector and agent-target distance predicted memory errors, including reminder use and non-target visits, in no-distraction trials. d). Time courses of object location, updated object vector and initial object vector strength during 2s before delay 2 and delay 2 period. b). Mean representational strength during delay 2. c). Trial-level GLMMs testing whether representational strength of object location, updated object vector, initial object vector and agent-target distance predicted memory errors, including reminder use and non-target visits, in distraction trials. In a & d, shaded areas in line plots indicate standard errors. Horizontal colored lines indicate above chance level representation strength. In b-f, error bars represent 95% confidence intervals. n.s indicates p > .05; * indicates p < .05. *** indicates p < .001.

## Relating representational strength to memory errors in memory load 2 trials

We tested if representational strength of maintained spatial information predicted trial-level memory errors, including reminder use and non-target visits. For no-distraction trials, none of the predictors significantly predicted whether the reminder was used in the trial (**SFig 5a**, agent-to-proximal target distance: β = 0.26, *CI* = [-0.16, 0.67], *BF10* = 1.16; agent-to-distal target distance: β = 0.11, *CI* = [-0.30, 0.51], *BF10* = 0.62; proximal object location: β = -0.31, *CI* = [-0.93, 0.31], *BF10* = 1.23; distal object location: β = -0.88, *CI* = [-1.82, 0.08], *BF10* = 5.68; proximal object vector: β = -0.07, *CI* = [-0.56, 0.43], *BF10* = 0.67; distal object vector: β = 0.34, *CI* = [-0.52, 1.17], *BF10* = 1.46; prospective object vector: β = 0.30, *CI* = [-0.53, 1.12], *BF10* = 1.34). For non-target visits, only the strength of distal object vector in delay 1 predicted fewer non-target visits (**SFig 5b**, β = -0.43, *CI* = [-0.75, -0.13], *BF10* = 16.58), but not for other predictors (agent-to-proximal target distance: β = 0.08, *CI* = [-0.05, 0.20], *BF10* = 0.33; agent-to-distal target distance: β = 0.12, *CI* = [-0.00, 0.24], *BF10* = 0.79; proximal object location: β = -0.10, *CI* = [- 0.29, 0.10], *BF10* = 0.39; proximal object vector: β = 0.01, *CI* = [-0.14, 0.16], *BF10* = 0.18; distal object vector: β = 0.24, *CI* = [-0.04, 0.52], *BF10* = 1.50; prospective objevt vector: β = 0.14, *CI* = [-0.13, 0.42], *BF10* = 0.58).

For distraction trials, as the agent-to-distal object distance increased, participants were more likely to use reminders (**SFig 5c**, β = 0.33, *CI* = [0.16, 0.51], *BF10* = 323.20), and visit non-target locations (**SFig 5d**), β = 0.19, *CI* = [0.09, 0.28], *BF10* = 219.87; As the agent-to-proximal object distance increased, participants were more likely to use reminders, β = 0.19, *CI* = [0.01, 0.36], *BF10* = 2.10. The strength of the prospective object vector was weakly associated with increased non-target visits (β = 0.20, *CI* = [0.02, 0.39], *BF10* = 2.42), and no other predictors reliably predicted reminder use (proximal object location: β = 0.14, *CI* = [-0.08, 0.35], *BF10* = 0.48; distal object location: β = 0.04, *CI* = [-0.36, 0.45], *BF10* = 0.69; proximal object vector: β = 0.15, *CI* = [-0.07, 0.38], *BF10* = 0.67; distal object vector: β = -0.18, *CI* = [-0.57, 0.19], *BF10* = 0.69; prospective object vector: β = 0.11, *CI* = [-0.23, 0.44], *BF10* = 0.51) or non-target visits (agent-to-proximal object distance: β = -0.01, *CI* = [-0.10, 0.08], *BF10* = 0.12; proximal object location: β = -0.02, *CI* = [-0.14, 0.10], *BF10* = 0.17; distal object location: β = -0.09, *CI* = [-0.31, 0.13], *BF10* = 0.40; proximal object vector: β = 0.10, *CI* = [-0.01, 0.22], *BF10* = 0.66; distal object vector: β = -0.13, *CI* = [-0.35, 0.07], *BF10* = 0.60).

**SFig. 5.**
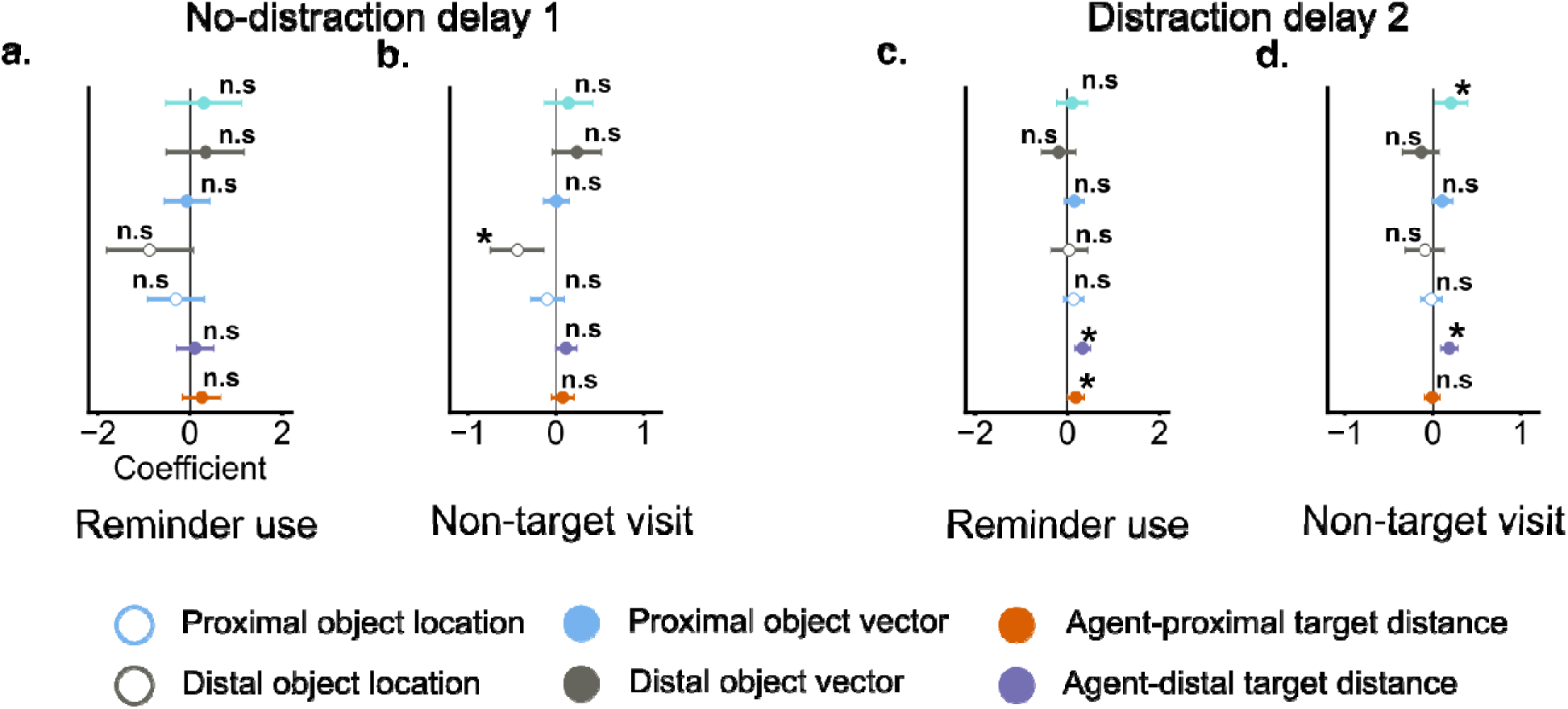
Linking object location and object vector coding in memory load 2 trials to behavioral memory errors. a&b). GLMM results linking representational strength during the delay 1 window to trial-level reminder use and non-target visits in no-distraction trials. c&d). GLMM results linking representational strength during the delay 2 window to trial-level reminder use and non-target visits in distraction trials. error bars represent 95% confidence intervals. n.s indicates p > .05; * indicates p < .05.

## Control analyses of attentional bias during delay windows

We repeated the analysis of fixation proportions after resampling each participant’s eye-tracking data to their lowest observed empirical sampling rate. The result patterns suggest that our fixation results were robust to variability in the empirical sampling rate.

During encoding and delay 1, fixations were more likely to fall near the proximal than the distal target after controlling for differences in agent-to-target distance (**SFig 6a**). In delay 1, the mean proportion of fixations on the proximal target was significantly greater than that on the distal object (**SFig 6b**), *t*(23) = 3.00, *p* = .006, *d* = 0.69, *BF10* = 7.08. During the distraction and delay 2 periods. Fixations were again more likely to fall near the proximal than the distal target (**SFig 6c**). During delay 2, the mean proportion of fixations on the proximal target was significantly greater than that on the distal target (**SFig 6d**), *t*(23) = 4.49, *p* <.001, *d* = 1.39, *BF10* = 173.39.

We next compared fixation patterns between distraction trials in which the optimal collection order changed following distraction (switch trials) and trials in which it remained unchanged (stay trials). This analysis tested whether plan updating was accompanied by corresponding shifts in fixation bias (**SFig 6e**). A repeated-measures ANOVA with factors of factors of time (pre-distraction vs. post-distraction), trial type (stay vs. switch), and item (initially proximal object, initially vs. initially distal object) revealed a significant three way interaction, *F*(1, 23) = 11.77, *p* = .002, 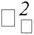 = .34. Followed-up ANOVAs revealed a significant interaction between item and time for switch trials, *F*(1, 23) = 13.33, *p* = .001, 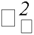 = .16, but not stay trials, *F*(1, 23) = 1.46, *p* = .239, 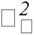 = .02. On switch trials, the initially proximal item became the distal item after distraction, and the proportions of fixations landing on it decreased [*t*(23) = -3.05, *p* = .023, *d* = 0.91, *BF10* = 7.76]. In contrast, the initially distal item became the proximal item after distraction, and proportions of fixations landing on it increased, though not statistically significant, *t*(23) = 2.73, *p* = .047, *d* = 0.83, *BF10* = 4.19. On stay trials, the initially distal item became the proximal item after distraction, and proportions of fixations landing on it decreased, *t*(23) = -2.83, *p* = .038, *d* = 0.83, *BF10* = 5.03. No significant difference between pre-distraction and post-distraction was found for initially proximal objects, *t*(23) = -0.27, *p* = .998, *BF10* = 0.22. Together, those result patterns suggest that our fixation results were robust to sampling rate.

**SFig. 6.**
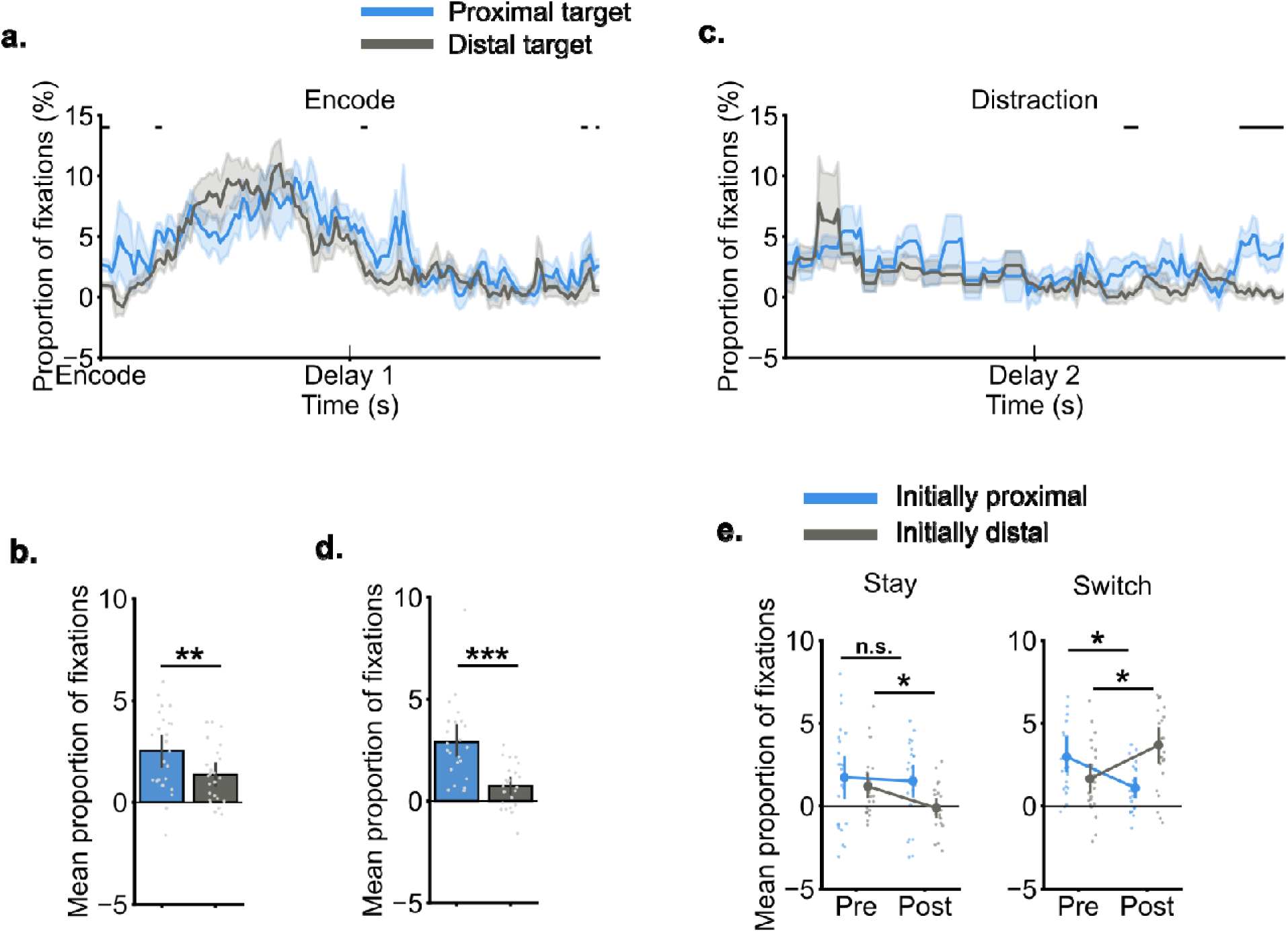
Fixation results in control analyses. a). Fixations on object locations as a function of time during encoding and the first delay window. b). During the delay following encoding, more fixations landed on proximal objects than on distal objects. c). Fixations on object locations as a function of time during the last two seconds of the distraction phase and the second delay window. d). During the delay following distractions, more fixations landed on updated proximal objects than on updated distal objects. e). Changes in proportions of fixations on items pre-distraction and post-distraction in stay and switch trials. Error bars indicate 95% confidence intervals. n.s indicates p > .05; * indicates p < .05. *** indicates p < .001. In a and c, horizontal colored lines indicate significant differences between the close and the far objects. The shaded area represents standard errors.

## Additional analyses of gaze trajectories during the distraction phase

To examine the functional relevance of gaze patterns, we used the rate of gaze patterns in a trial to predict whether that trial ended with showing optimal behavior in distraction trials (**Sfig 7a**). No gaze patterns reliably predicted optimal behavior (encode proximal: β = 0.09, *CI* = [-0.02, 0.22], *BF10* = 0.47; encode distal: β = 0.01, *CI* = [-0.10, 0.13], *BF10* = 0.14; optimal forward: β = 0.09, *CI* = [-0.04, 0.25], *BF10* = 0.33; suboptimal forward: β = -0.04, *CI* = [-0.14, 0.07], *BF10* = 0.17; optimal backward: β = 0.01, *CI* = [-0.11, 0.12], *BF10* = 0.14; suboptimal backward: β = -0.01, *CI* = [-0.11, 0.11], *BF10* = 0.14).

We conducted control analyses to assess whether gaze pattern results were robust to variability in the empirical sampling rate (**Sfig 7b**). Specifically, we repeated the analyses after resampling each participant’s eye-tracking data to their lowest observed empirical sampling rate. The result patterns suggest that our fixation results were robust to variability in the empirical sampling rate.

A repeated-measures ANOVA revealed a significant main effect of gaze pattern, *F*(5, 115) = 27.72, *p* < .001, 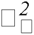 = .43. At this reduced sampling rate, we replicated the main finding that backward sweeps aligned with optimal plans occurred more frequently than those aligned with suboptimal plans, *t*(23) = 3.53, *p* = .002, *d* = 0.71, *BF10* = 21.03. No significant difference between optimal and suboptimal forward sweeps was found, *t*(23) = -1.23, *p* = .232, *BF10* = 0.42. Gaze patterns consistent with the independent encoding of the proximal target occurred more frequently than those associated with the distal target, *t*(23) = 6.35, *p* < .001, *d* = 1.78, *BF10* = 1.1 × 10^4^.

We compared the occurrence rates of each gaze pattern between stay and switch trials. A repeated-measures ANOVA with variables of trial type (stay vs. switch) and gaze pattern types revealed a significant interaction, *F*(5, 115) = 4.55, *p* < .001, 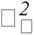 = .17. Follow-up comparisons showed that only backward sweeps occurred more frequently on switch than on stay trials (**Sfig 7c**), *t*(24) = 5.21, *p* < .001, *d* = 0.94, *BF10* = 871.64. No other gaze patterns differed significantly between stay and switch trials (*ps* > 0.3).

**SFig. 7.**
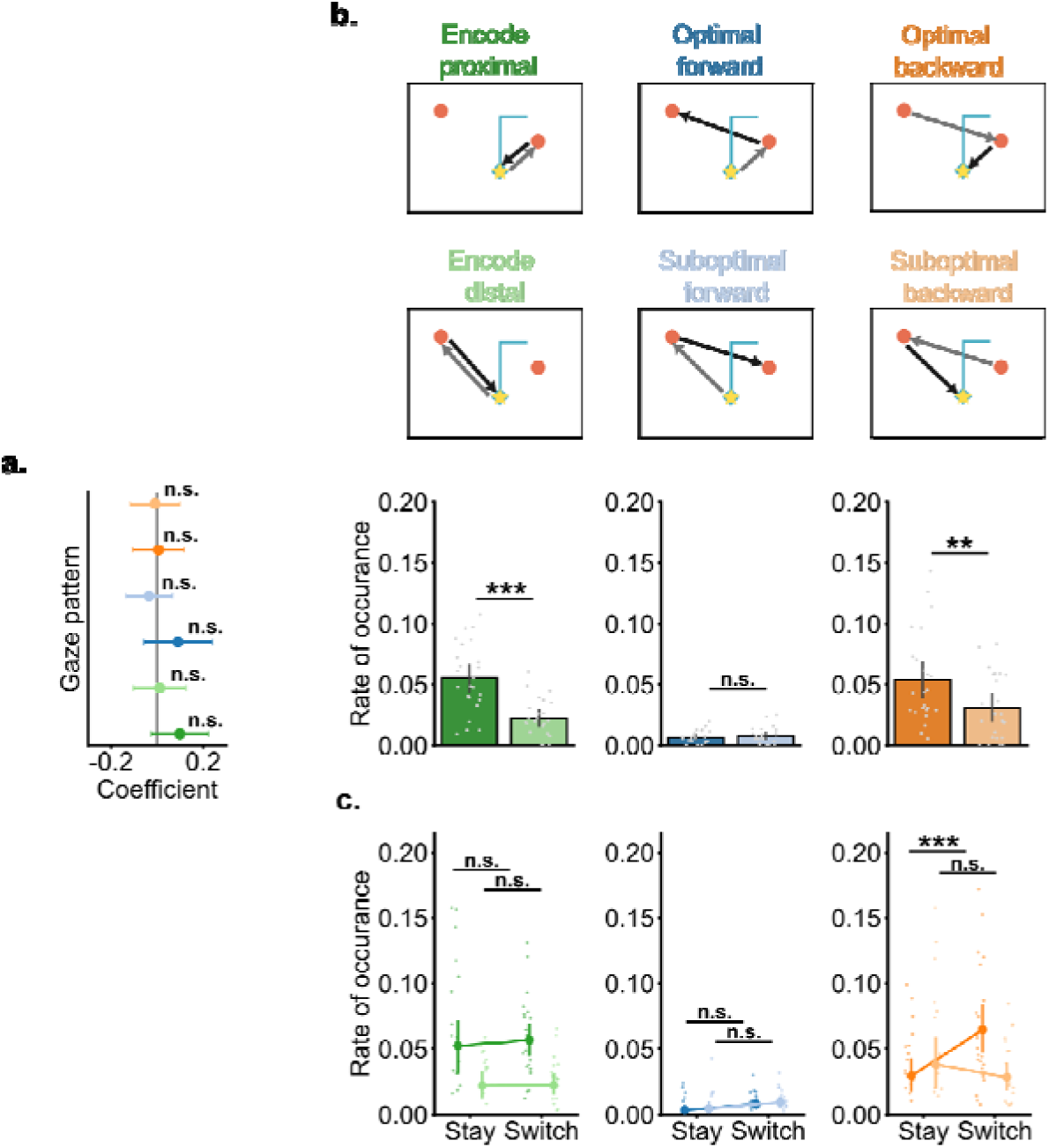
Additional analyses of gaze trajectories during the distraction window. a). Trial-level GLMM relating the rate of each gaze pattern to whether participants subsequently selected the optimal collection order in distraction trials. b). Rate of each gaze pattern after resampling the eye-tracking data to a reduced sampling rate. Error bars represent 95% confidence intervals. c). Rate of occurrence for each gaze pattern for stay and switch trials. n.s indicates p > .05; ** indicates p < .01; *** indicates p < .001.

